# A next generation approach to species delimitation reveals the role of hybridization in a cryptic species complex of corals

**DOI:** 10.1101/523936

**Authors:** Andrea M. Quattrini, Tiana Wu, Keryea Soong, Ming-Shiou Jeng, Yehuda Benayahu, Catherine S. McFadden

## Abstract

**Background:** Our ability to investigate processes shaping the evolutionary diversification of corals (Cnidaria: Anthozoa) is limited by a lack of understanding of species boundaries. Discerning species has been challenging due to a multitude of factors, including homoplasious and plastic morphological characters and the use of molecular markers that are either not informative or have not completely sorted. Hybridization can also blur species boundaries by leading to incongruence between morphology and genetics. We used traditional DNA barcoding and restriction-site associated DNA sequencing combined with coalescence-based and allele-frequency methods to elucidate species boundaries and simultaneously examine the potential role of hybridization in a speciose genus of octocoral, *Sinularia*.

**Results:** Species delimitations using two widely used DNA barcode markers, *mtMutS* and 28S rDNA, were incongruent with one another and with the morphospecies identifications, likely due to incomplete lineage sorting. In contrast, 12 of the 15 morphospecies examined formed well-supported monophyletic clades in both concatenated RAxML phylogenies and SNAPP species trees of >6,000 RADSeq loci. DAPC and Structure analyses also supported morphospecies assignments, but indicated the potential for two additional cryptic species. Three morphologically distinct species pairs could not, however, be distinguished genetically. ABBA-BABA tests demonstrated significant admixture between some of those species, suggesting that hybridization may confound species delimitation in *Sinularia*.

**Conclusions:** A genomic approach can help to guide species delimitation while simultaneously elucidating the processes generating diversity in corals. Results support the hypothesis that hybridization is an important mechanism in the evolution of Anthozoa, including octocorals, and future research should examine the contribution of this mechanism in generating diversity across the coral tree of life.

## Background

The ability to delimit species is fundamental to the accurate assessment of biodiversity and biogeography, information that is essential for studying their biology as well as for implementing conservation policies. Yet this task is not trivial, as species are often difficult to discriminate for a multitude of reasons. Morphological traits have traditionally been used in classical taxonomy; however, use of characters that might not be diagnostic or are homoplasious can confound the interpretation of species boundaries. Cryptic species, particularly those that occur in sympatry, and species that have arisen via hybridization and introgression are often challenging to discriminate without genetic, ecological or behavioral data. DNA barcoding of mitochondrial genes has proven useful in many species groups (Hebert et al., 2004a, b), but incomplete lineage sorting and past hybridization events complicate species delimitation based on mitochondrial data alone, particularly in recently diverged taxa (Hickerson et al., 2006; Johnston et al., 2017; McFadden et al., 2017). In addition, mitochondrial markers reflect the history of maternal lineages, which are often incongruent with the species history (Currat et al., 2008; Rheindt & Edwards, 2011). The increased resolution of genomic data can potentially disentangle some of these issues, facilitating species delimitation while simultaneously furthering our understanding of processes that generate biodiversity (Sukumaran & Knowles, 2017). Moreover, such an approach may also provide a better evaluation of morphological traits and insights into their congruence with genetic data.

In sessile marine invertebrates, such as corals, congeners often occur in sympatry and occupy similar ecological niches and reef zones. These ecological characteristics combined with reproductive modes may lead to increased rates of hybridization among close relatives. Broadcast-spawning species that occur in sympatry often participate in synchronous, mass-spawning reproduction events (Babcock et al. 1986; Harrison et al., 1984; Kahng et al. 2011; Richmond & Hunter, 1990). Unless there are prezygotic mechanisms to reproductive isolation, such as gametic incompatibility or asynchronous spawning times, there may be numerous opportunities for hybridization to occur (van Oppen et al., 2002; Willis et al., 1997, 2006). In fact, laboratory crossings of sympatric congeners have produced viable hybrid offspring in several species (Slattery et al., 2008; Willis et al., 1997). Hybridization followed by reticulate evolution has been suggested to be an important mechanism generating the species diversity observed in some groups of corals (Combosch & Vollmer, 2015; Diekmann et al., 2001; Frade et al., 2010; Hatta et al., 1999; McFadden & Hutchinson, 2004; Miller & van Oppen, 2003; Richardson et al., 2008; van Oppen et al., 2001, 2002; Willis et al., 2006). Vollmer and Palumbi (2002), however, suggested that hybridization could yield distinct, new morphotypes that may be reproductively inviable or subject to hybrid breakdown. It is clear that further investigation is needed to determine the potential contributions of hybridization to speciation and morphological innovation in corals.

One particularly speciose group of octocorals (Cnidaria: Anthozoa: Octocorallia) is the genus *Sinularia* May, 1898. This zooxanthellate genus includes approximately 175 valid species (WoRMS Editorial Board 2018), 47 of them described just in the last 25 years. They are diverse and abundant throughout the Indo-Pacific, and biodiversity surveys of shallow-water coral reef communities typically report more than 15 co-occurring species of *Sinularia*, with as many as 38 species recorded at some locations (Manuputty & Ofwegen, 2007; Ofwegen, 2002, 2008). *Sinularia* species are typically most abundant on reef flats and shallow slopes, where single- or multi-species assemblages may dominate the reef substrate (Benayahu, 1995; Benayahu & Loya, 1977; Dinesen, 1983; Fabricius, 1998; Tursch & Tursch, 1982). *Sinularia* can also play an active role in reef biogenesis through deposition of spiculite formed by the cementation of layers of calcitic sclerites (Jeng et al., 2011; Shoham et al. in press). In addition, many *Sinularia* species produce secondary metabolites used for allelopathy and predator deterrence (Slattery et al., 1999, 2001; van Alstyne et al., 1994; Wylie & Paul, 1989), making the genus a rich and diverse source of bioactive natural products (e.g., Blunt et al., 2016).

Because of the dominance and importance of *Sinularia* species across a wide depth gradient (Shoham & Benayahu, 2017) as well as their susceptibility to bleaching-induced mortality (Bruno et al., 2001; Fabricius, 1999; Goulet et al., 2006; Marshall & Baird, 2000), it is of a great interest to better understand their ecology and function on the reefs. Ecological studies, however, are often hampered by the uncertainty of species identifications (McFadden et al., 2009). Classical taxonomy of *Sinularia* species is based primarily on morphological features of the colony and the shape and dimension of sclerites (microscopic calcitic skeletal elements) found in different parts of the colony (Fabricius & Alderslade, 2001; McFadden et al., 2009; Verseveldt, 1980). Separation of species using these characters can be subjective, as the complex morphologies of both colonies and sclerites are rarely quantified (Aratake et al., 2012; Carlo et al., 2011). There is also a potential contribution of environmental plasticity to the morphological variation observed, as has been documented in other octocorals (Kim et al., 2004; Rowley et al., 2015; Sánchez et al., 2007).

The application of molecular systematic and DNA barcoding approaches to the study of species boundaries in *Sinularia* have been only partially successful (McFadden et al., 2009). While molecular approaches have revealed that some well-known morphospecies comprise cryptic species complexes (Ofwegen et al., 2013, 2016), it is also the case that numerous morphologically distinct *Sinularia* species share identical haplotypes at barcoding loci (Benayahu et al., 2018; McFadden et al., 2009, 2014). Because mitochondrial genes evolve slowly in Anthozoa (Huang et al., 2008; Shearer & Coffroth, 2008), these markers often simply lack the resolution to distinguish recently diverged species (McFadden et al., 2011, 2014). As a result, it is often not possible to conclude with certainty whether morphologically distinct individuals that share identical DNA barcodes represent different octocoral species or morphological variants of a single species. In addition, the reported ability of some species of *Sinularia* to hybridize in the laboratory (Slattery et al., 2008), raises the possibility that naturally occurring hybridization events could contribute to the observed morphological diversity of this genus, as has been suggested for stony corals (Richards et al., 2008). The true identity of some *Sinularia* species remains uncertain, and our ability to explore the evolutionary processes leading to diversification of this hyperdiverse lineage are limited by a lack of understanding of species boundaries. In fact, we have a limited understanding of species boundaries for numerous groups of recently diverged corals (e.g., Johnston et al., 2017; McFadden et al., 2017; van Oppen et al., 2000).

Octocoral biodiversity surveys conducted recently at Dongsha Atoll, Taiwan, recorded 27 nominal morphospecies of *Sinularia* inhabiting the reef slope down to a depth of 20 m (Benayahu et al., 2018), most of them belonging to the speciose clades “4” and “5C” (McFadden et al., 2009). These two clades include several subclades each characterized by a different suite of morphological characters whose diagnosis is quite confusing (McFadden et al., 2009). While most of these morphospecies could be distinguished using a character-based mitochondrial gene barcode (*mtMutS*), five distinct morphospecies in clade 5C shared identical haplotypes, and several morphospecies in both clades were represented by more than one haplotype (Benayahu et al., 2018). These morphospecies exemplify a problem common to many corals and raise the following questions: (1) do the observed morphological differences reflect boundaries between species whose mitochondrial haplotypes have not yet diverged or coalesced, or do these differences reflect intraspecific variation? and (2) might the sharing of mitochondrial haplotypes among distinct morphotypes reflect ongoing or past hybridization events?

To further explore these questions and to elucidate species boundaries in the *Sinularia* that co-occur on Dongsha Atoll, we have (1) sequenced an additional, nuclear marker (28S rDNA) that has been shown to be comparable to *mtMutS* as a species-specific barcode for *Sinularia* and other octocoral taxa (McFadden et al., 2014); and (2) sequenced restriction-site associated DNA (RADseq) to identify SNPs for multilocus species delimitation analyses using allele-frequency and coalescence-based approaches. Expanding upon a recent study of co-occurring *Sinularia* species at Dongsha Atoll (Benayahu et al., 2018), we validate morphospecies identifications using a genomic approach, and provide insight into the possible role of hybridization in the evolution of the genus.

## Results

### Species Delimitation using DNA Barcodes

Neither the *mtMutS* (735 bp) nor the *28S rDNA* (764 bp) barcoding marker delimited all morphospecies of *Sinularia* when considered separately (Fig. 2). Phylogenetic relationships among morphospecies were poorly resolved with low support values and few reciprocally monophyletic groups, especially in clade 5C (Suppl. Fig. 1). Based on a 0.3% genetic distance threshold, the *mtMutS* barcode identified six molecular operational taxonomic units (MOTUs) among the four morphospecies belonging to clade 4, splitting *S. tumulosa* and *S. ceramensis* into two MOTUs each (Fig. 2a). In contrast, *28S rDNA* (0.3% threshold) delimited only four MOTUs within clade 4, each of them congruent with morphospecies identifications. The only exception was *S. verruca* (R41341) from Palau, included as a taxonomic reference, whose *mtMutS* and *28S* sequences were identical to those of *S. tumulosa*.

**Figure 1.**
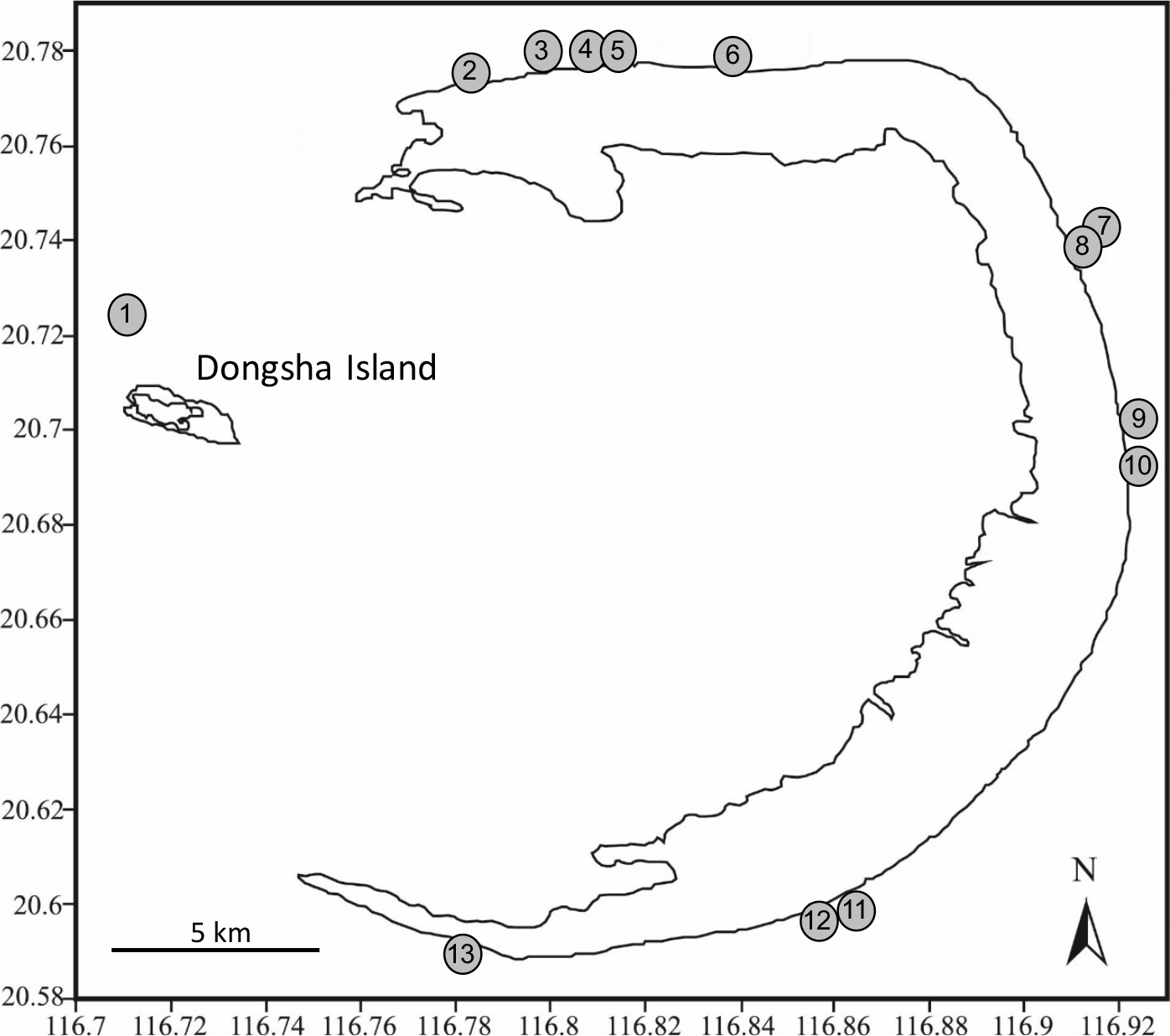
Map of Dongsha Atoll Marine National Park, Taiwan. Collection sites indicated by numbered circles. Adapted from Benayahu et al. (2018).

**Figure 2.**
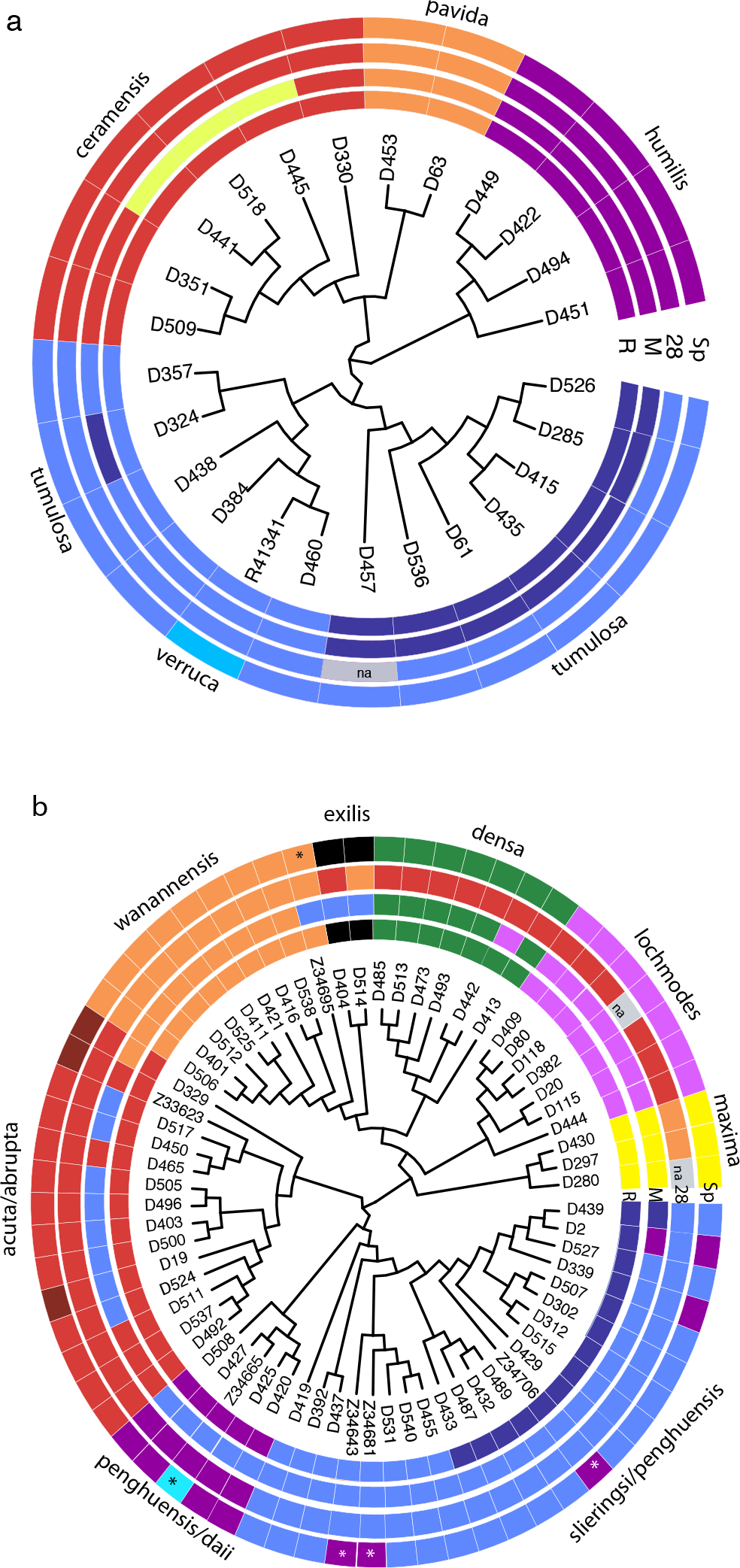
Maximum likelihood cladograms for Dongsha Atoll *Sinularia* a) clade 4 and b) clade 5C. Each colored cell denotes an individual’s species assignment based on RAD data (R), a morphospecies assignment (Sp), or a molecular operational taxonomic unit (MOTUs) based on *mtMutS* (M) and 28S rDNA (28). Colors match RAD clades in Figures 3 and 4. *=holotype and paratype. Morphospecies names are also included. See also Suppl. Table 1 for clade and MOTU assignments. Gray cells (‘na’) denote missing data.

Among the clade 5C morphospecies, *mtMutS* delineated eight MOTUs (Fig. 2b). With just two exceptions (D442, Z34695), *mtMutS* differentiated *S. maxima*, *S. wanannensis*, *S. lochmodes* and *S. densa* from all other morphospecies (Fig. 2b). A majority of the colonies identified as *S. penghuensis* and *S. slieringsi*, both individuals of *S. exilis*, and seven of eleven *S. acuta*, however, belonged to a single MOTU, while the remaining four individuals of *S. acuta* were assigned to a separate MOTU. Two individuals with divergent haplotypes (*S. penghuensis* D002 and *S. slieringsi* D439) were each assigned to unique MOTUs.

**Table 1.** Morphospecies of *Sinularia* included in RADseq analysis. Clade corresponds to designations in McFadden et al., (2009). N: number of specimens sequenced. #Sites: number of different dive sites at Dongsha Atoll at which a species was collected.

| Species | Authority | Clade | N | #Sites | Depth<br>(m) |
| --- | --- | --- | --- | --- | --- |
| *S. abrupta* | Tixier-Durivault, 1970 | 5C | 3 | 2 | 8–13 |
| *S. acuta* | Manuputty & Ofwegen, 2007 | 5C | 11 | 6 | 4–14 |
| *S. ceramensis* | Verseveldt, 1977 | 4D | 6 | 5 | 3–13 |
| *S. daii* * | Benayahu & Ofwegen, 2011 | 5C | 1 | - | --- |
| *S. densa* | (Whitelegge, 1897) | 5C | 6 | 4 | 3–14 |
| *S. exilis* | Tixier-Durivault, 1970 | 5C | 2 | 2 | 5–14 |
| *S. humilis* | Ofwegen, 2008 | 4B | 4 | 3 | 3–21 |
| *S. lochmodes* | Kolonko, 1926 | 5C | 8 | 6 | 4–14 |
| *S. maxima* | Verseveldt, 1971 | 5C | 3 | 2 | 3–17 |
| *S. pavida* | Tixier-Durivault, 1970 | 4D | 2 | 2 | 4–5 |
| *S. penghuensis* | Ofwegen & Benayahu, 2012 | 5C | 8 | 3 | 3–21 |
| *S. slieringsi* | Ofwegen & Vennam, 1994 | 5C | 17 | 6 | 3–21 |
| *S. tumulosa* | Ofwegen, 2008 | 4D | 12 | 8 | 3–17 |
| *S. verruca* * | Ofwegen, 2008 | 4D | 1 | - | --- |
| *S. wanannensis* | Ofwegen & Benayahu, 2012 | 5C | 9 | 5 | 5–21 |

\*reference species not collected at Dongsha

In contrast to *mtMutS*, *28S* delineated only four MOTUs within clade 5C (Fig. 2b). *S. penghuensis* and *S. slieringsi*, which were not separated using *mtMutS*, were separated into two distinct MOTUs; four of the *S. penghuensis* colonies belonged to a MOTU that was well separated from all others, while the other five—including the holotype (Z34706) and both paratypes—shared identical genotypes with *S. slieringsi*. A third MOTU included *S. acuta*, *S*. *densa* and *S. lochmodes* along with *S. abrupta* and one *S. exilis*. The fourth MOTU included all individuals of *S. maxima* and *S. wanannensis* plus the second *S. exilis*.

Several other colonies also had *mtMutS* or *28S* genotypes that were not consistent with their morphospecies identity. The holotype of *S. daii* from Penghu (Z34665) had *mtMutS* and *28S* sequences identical to those of some *S. penghuensis*. Among the three colonies identified as *S. abrupta*, D329 had a *mtMutS* sequence matching *S. wanannensis* but a *28S* sequence matching *S. acuta*; Z33623 (from Penghu) had a *mtMutS* sequence matching *S. acuta* but its *28S* matched *S. densa*; and D019 was identical to *S. acuta* at both loci. The holotype of *S. wanannensis* from Penghu (Z34695) had a *28S* sequence consistent with other individuals of that species, but shared a *mtMutS* haplotype with *S. penghuensis* and *S. slieringsi* (Fig. 2b).

### RADSeq Data Statistics

A total of 289,373,374 reads were obtained for 95 *Sinularia* samples. After trimming in both Stacks and pyRAD, 86% of reads were retained (247,873,622). The mean number of reads per individual was 2,609,196 ± 627,314. For each clade, the number of loci and the number of SNPs obtained increased considerably when the number of shared heterozygous sites (*p)* was increased and both the clustering threshold (*c*) and individual occupancy per locus (m) were decreased (Table 2). Notably, a substantial increase in both the number of loci and SNPs obtained occurred when *p* was set to 0.25 at a clustering threshold (*c*) of 0.85. The number of loci obtained ranged from 73 to 28,179 for clade 4 and 115 to 23,946 for clade 5C depending upon parameters used in pyRAD analyses (Table 2). The number of variable SNPs obtained ranged from 382 to 251,615 for clade 4 and 885 to 329,837 for clade 5C (Table 2).

**Table 2.** Dongsha Atoll *Sinularia*. Loci and SNP summary statistics of pyRAD simulations at two different clustering thresholds (*c*), three different levels of taxon occupancy per locus (*m)*, and two different levels of shared polymorphic sites *(p)*. Minimum and maximum loci obtained for one individual are included as well as total loci obtained across all individuals. Total number of variable SNPs (Var), parsimony informative SNPs (PI), and unlinked bi-allelic SNPs (BI) are also included.

|  |  |  |  | Number of Loci |  |  |  | Number of SNPs |  |  |  |  |  |
| --- | --- | --- | --- | --- | --- | --- | --- | --- | --- | --- | --- | --- | --- |
| (m) | (c) | Min | Max | Total | Min | Max | Total | Var. | PI | BI | Var. | PI | BI |
| Clade 4 |  |  | *p10* |  |  | *p25* |  |  | *p10* |  |  | *p25* |  |
| 1.0 | 0.90 | 73 | 73 | 73 | 154 | 154 | 154 | 382 | 78 | 63 | 1,358 | 617 | 144 |
|  | 0.85 | 93 | 93 | 93 | 185 | 185 | 185 | 563 | 255 | 83 | 1,675 | 778 | 175 |
| 0.75 | 0.90 | 1,570 | 2,391 | 2,552 | 3,139 | 5,331 | 5,515 | 17,690 | 10,505 | 2,448 | 48,301 | 28,211 | 5,411 |
|  | 0.85 | 1,869 | 2,869 | 2,958 | 3,791 | 6,113 | 6,343 | 22,936 | 14,026 | 2,851 | 60,232 | 36,467 | 6,236 |
| 0.50 | 0.90 | 3,391 | 9,742 | 10,813 | 6,923 | 24,178 | 26,861 | 70,048 | 38,239 | 10,490 | 214,870 | 121,983 | 26,532 |
|  | 0.85 | 4,347 | 10,341 | 11,540 | 8,693 | 25,221 | 28,179 | 86,607 | 49,558 | 11,225 | 251,615 | 147,779 | 27,860 |
| Clade 5C |  |  |  |  |  |  |  |  |  |  |  |  |  |
| 1.0 | 0.90 | 115 | 115 | 115 | 143 | 143 | 143 | 885 | 212 | 104 | 1,222 | 356 | 132 |
|  | 0.85 | 123 | 123 | 123 | 154 | 154 | 154 | 908 | 229 | 109 | 1,325 | 385 | 140 |
| 0.75 | 0.90 | 2,500 | 4,174 | 4,306 | 4,060 | 6,675 | 7,083 | 52,325 | 31,844 | 4,276 | 91,810 | 57,263 | 7,051 |
|  | 0.85 | 2,781 | 4,793 | 4,968 | 4,491 | 7,771 | 8,060 | 62,057 | 38,113 | 4,930 | 106,548 | 66,726 | 8,022 |
| 0.50 | 0.90 | 6,210 | 11,587 | 13,189 | 9,439 | 20,926 | 23,946 | 171,091 | 104,614 | 13,141 | 329,837 | 205,990 | 23,898 |
|  | 0.85 | 5,381 | 11,697 | 13,281 | 9,329 | 20,582 | 23,484 | 175,558 | 108,258 | 13,225 | 327,211 | 205,557 | 23,428 |

### Phylogenetic Inference and Species Delimitation

In contrast to the *mtMutS* and *28S* rDNA trees (Suppl. Fig. 1), a majority of the identified morphospecies formed well-supported monophyletic clades in both clade 4 and 5C phylogenies constructed with the *c* 0.85, *p* 0.25, and *m* 0.75 RADSeq datasets (clade 4: 6,343 loci, clade 5C: 8,060 loci; Figs. 3-4). The maximum clade credibility species trees produced from SNP data in the SNAPP analyses were largely congruent with the RAxML trees generated from concatenated data (Fig. 5). However, in clade 4, *S. pavida* and *S. ceramensis* were reciprocally monophyletic in the ML tree but not in the maximum clade credibility SNAPP species tree, although this relationship was evident in 30% of the alterative SNAPP tree topologies (in red, Fig. 5A). In clade 5C, *S. acuta* was sister to *S. penghuensis* and *S. slieringsi* in the ML tree, but not in the maximum clade credibility species tree, although this relationship was evident in 25% of the alternative SNAPP tree topologies (in red and green, Fig. 5B). For clade 4, 35% of the SNAPP trees obtained were alternative topologies to the maximum clade tree and 37% of trees had different topologies compared to the maximum clade credibility tree for clade 5C (Fig. 5).

**Figure 3.**
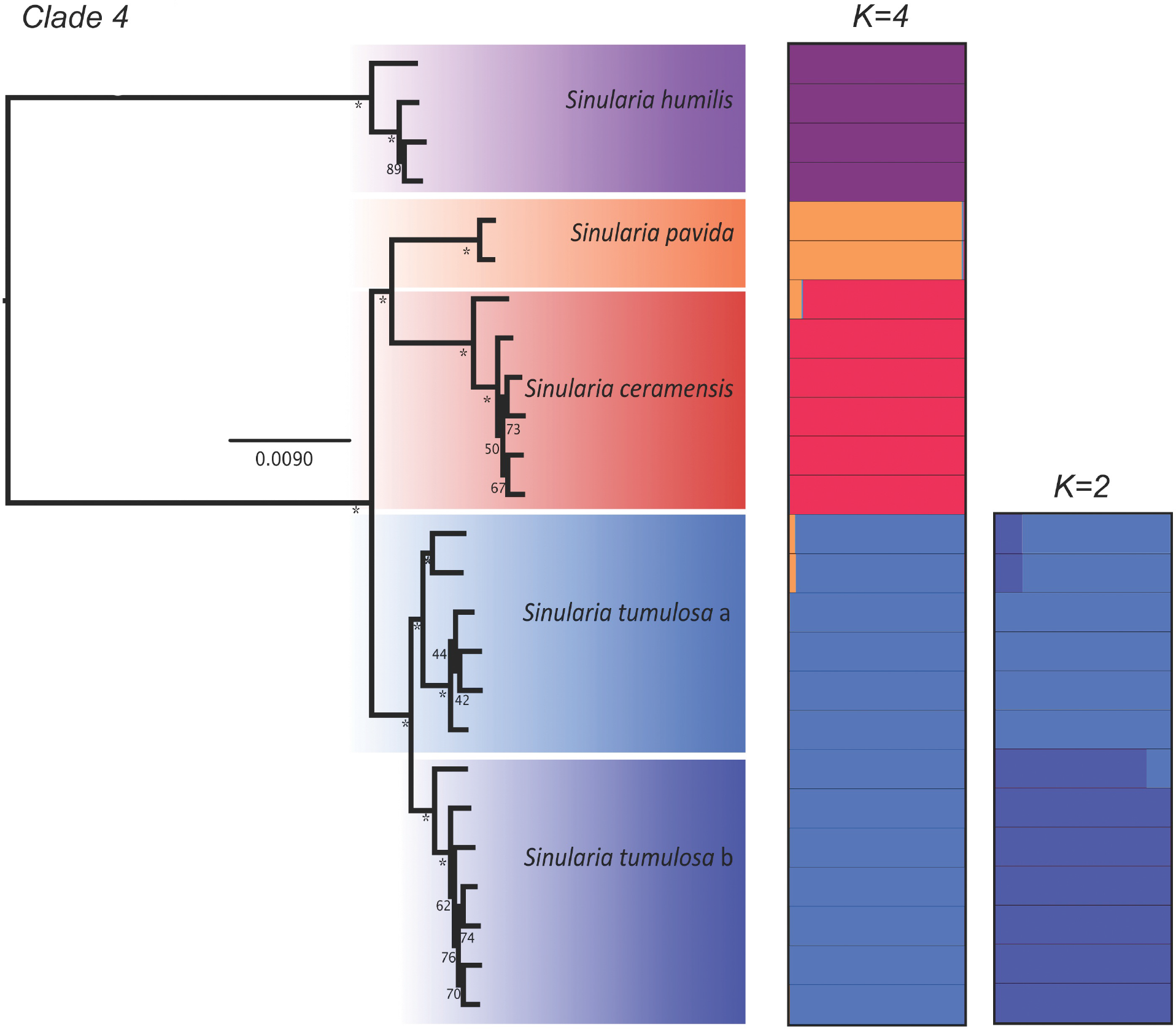
Maximum likelihood phylogeny of Dongsha Atoll *Sinularia* clade 4 constructed using RAxML rapid bootstrapping (200 b.s. replicates) on the concatenated c 0.85, m 0.75, p 0.25 locus dataset. * denotes 100% b.s. support. Distruct plots are included and show the probability of individual membership into different *K* (*K*=4 for clade4 and *K*=2 for the *S. tumulosa* group) clusters. Colors denote different species.

**Figure 4.**
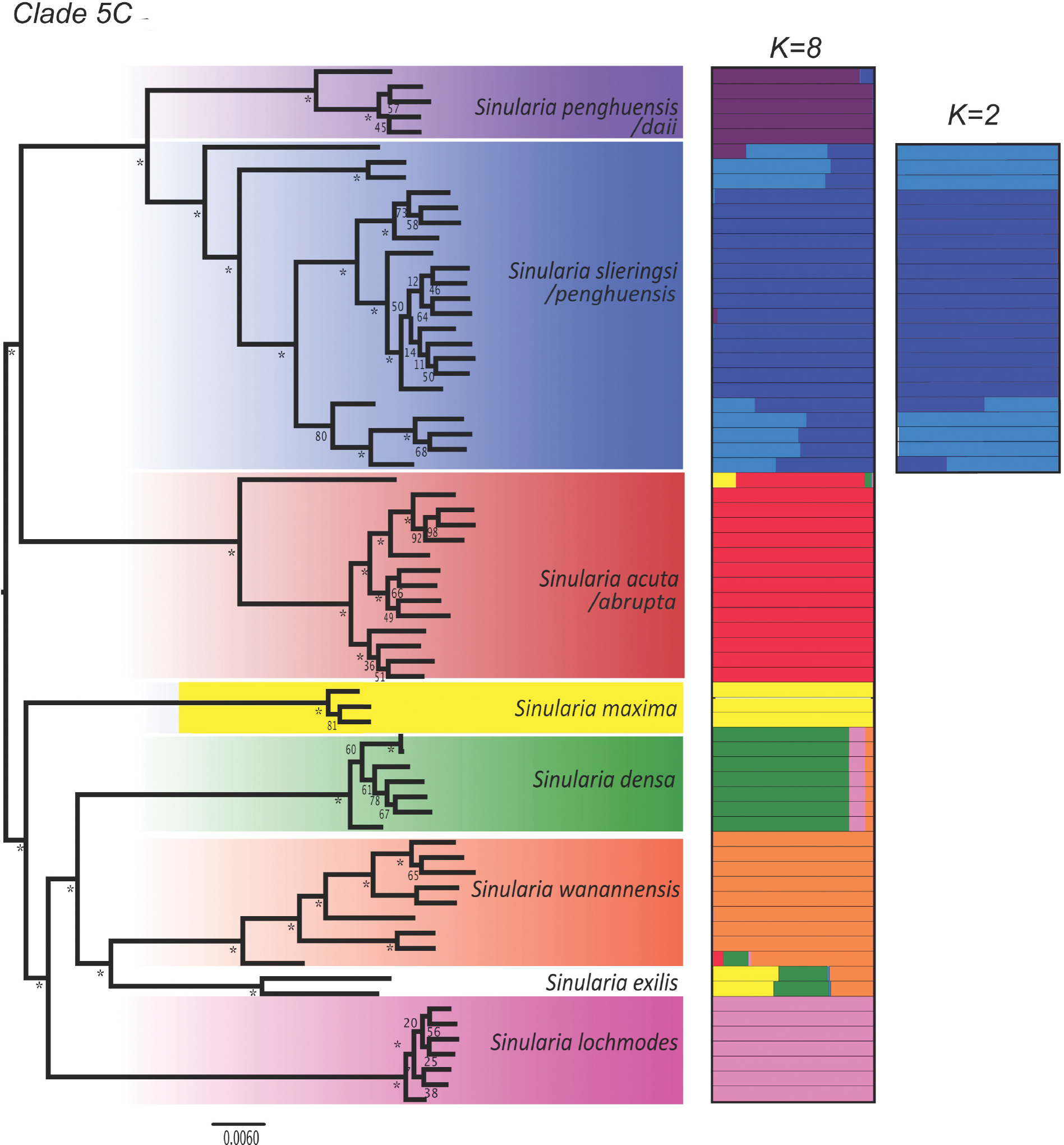
Maximum likelihood phylogeny of Dongsha Atoll *Sinularia* clade 5C constructed using RAxML rapid bootstrapping (200 b.s. replicates) on the concatenated c 0.85, m 0.75, p 0.25 locus dataset. * denotes 100% b.s. support. Distruct plots are included and show the probability of membership into different *K* (*K*=8 for clade 5C, and *K*=2 for the *S*. *slieringsi* group) clusters. Colors denote different species.

**Figure 5.**
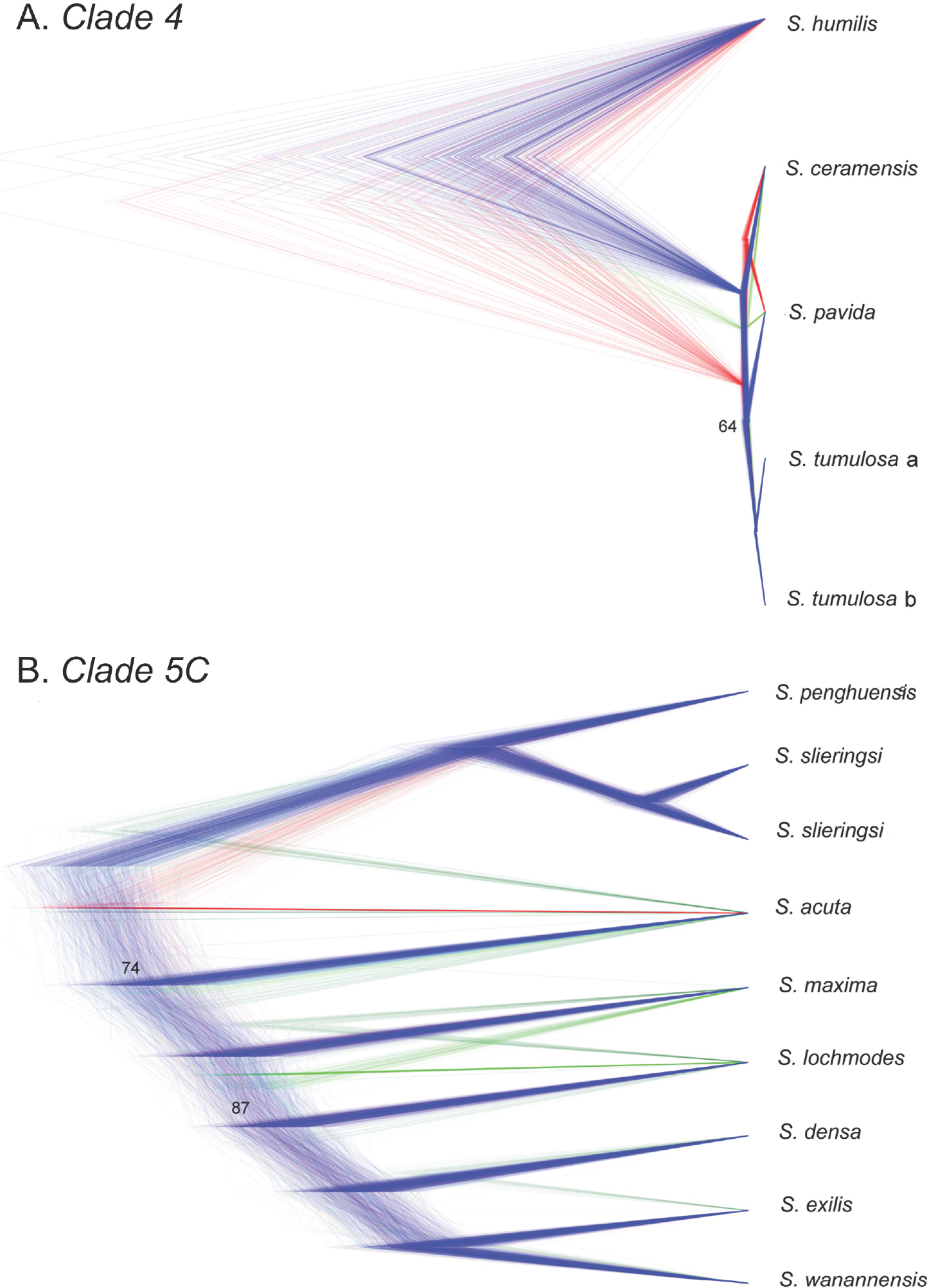
Species trees of Dongsha Atoll *Sinularia* clades 4 and 5C. Cloudograms illustrate the best species delimitation models (DAPC+1) for both clades inferred from bi-allelic SNP data [(A) clade 4: 6,236 SNPs and (B) clade 5C: 8,022 SNPs); *m*0.75 datasets] using SNAPP species tree analyses. The maximum clade credibility tree and congruent trees are in blue. Trees with different topologies are in red and green. Posterior probabilities at internal nodes >95% unless indicated.

### Clade 4

Species delimitation analyses agreed with the currently defined morphospecies in *Sinularia* clade 4 (*S. ceramensis*, *S. humilis, S. pavida*, and *S. tumulosa*), and with the four MOTUs identified by the *28S rDNA* barcoding marker (Fig. 2a). Consistent with the barcoding results, *S. verruca* was genetically indistinguishable from *S. tumulosa*. The optimal number of *K* clusters suggested by the DAPC analyses was four (BIC=120.4, Suppl. Fig. 2), and the DAPC plot revealed no overlap among these four distinct clusters (Fig. 6a). Further support for group assignment can be seen in the assignment plots, as all individuals were successfully re-assigned into their respective clusters (Suppl. Fig. 3). In addition, the Distruct plot clearly illustrated little to no admixture among these four species (Fig. 3). Upon further Structure analysis, little to no admixture was also revealed between two sub-clades of *S. tumulosa* (Fig. 3), suggesting that *S. tumulosa* might consist of two species. Other methods also support this result. First, following a one-species model (MLE=−1119), DAPC+1 was the second most likely (MLE=−1291) species model according to BFD* analyses (Table 3). The DAPC+1 model included species denoted by DAPC, plus two sub-clades of *S. tumulosa*. Second, most of the individuals of these two sub-clades formed two separate groupings in the DAPC plot, although there was some overlap among individuals (Fig. 6a). Finally, *S. tumulosa* was divided into two well-supported, reciprocally monophyletic clades (sp. a and b) in both concatenated and species tree phylogenies, which match the *mtMutS* results (Figs. 2a, 3 and 5a). It is possible that these represent two cryptic species.

**Table 3.**
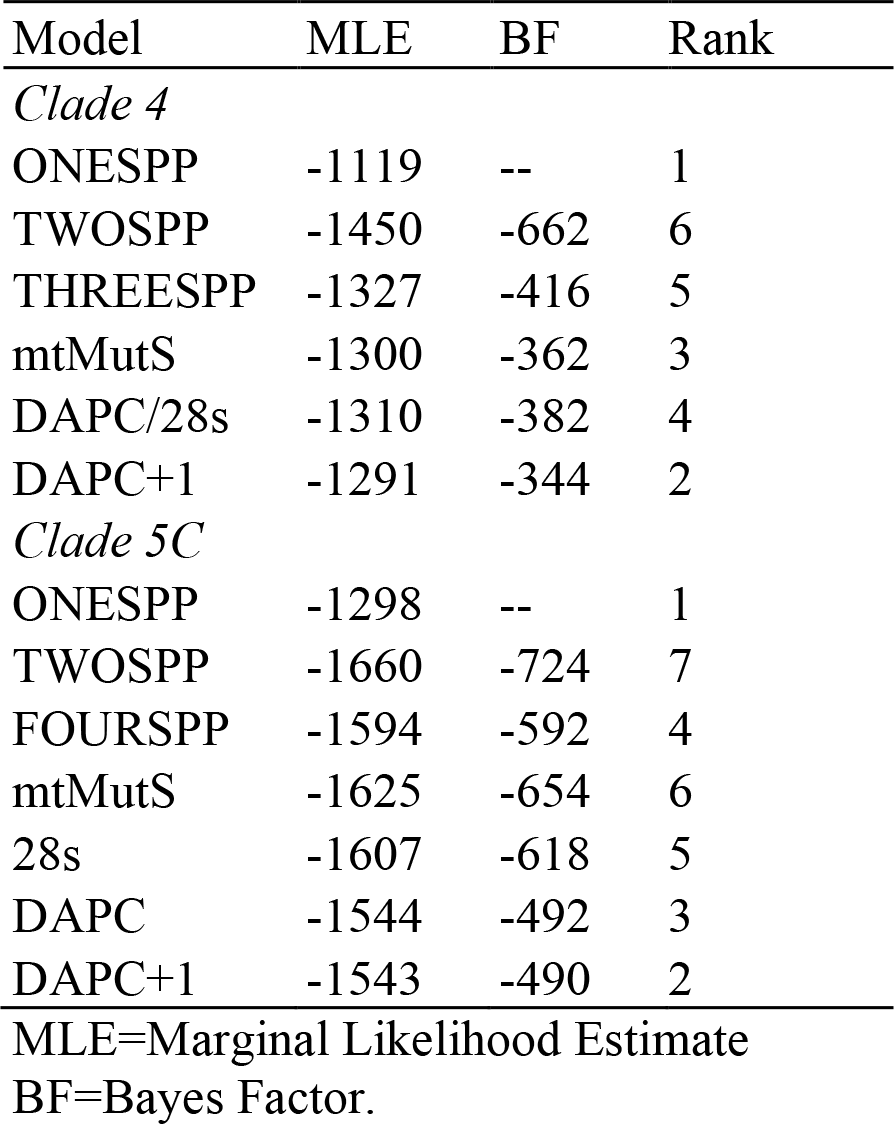
SNAPP results for different species delimitation models for Dongsha Atoll *Sinularia* clades 4 and 5C. Rank of most likely species model based on Bayes Factor delimitation is indicated.

| Model | MLE | BF | Rank |
| --- | --- | --- | --- |
| *Clade 4* |  |  |  |
| ONESPP | -1119 | -- | 1 |
| TWOSPP | -1450 | -662 | 6 |
| THREESPP | -1327 | -416 | 5 |
| mtMutS | -1300 | -362 | 3 |
| DAPC/28s | -1310 | -382 | 4 |
| DAPC+1 | -1291 | -344 | 2 |
| *Clade 5C* |  |  |  |
| ONESPP | -1298 | -- | 1 |
| TWOSPP | -1660 | -724 | 7 |
| FOURSPP | -1594 | -592 | 4 |
| mtMutS | -1625 | -654 | 6 |
| 28s | -1607 | -618 | 5 |
| DAPC | -1544 | -492 | 3 |
| DAPC+1 | -1543 | -490 | 2 |

MLE=Marginal Likelihood Estimate
BF=Bayes Factor.

**Figure 6.**
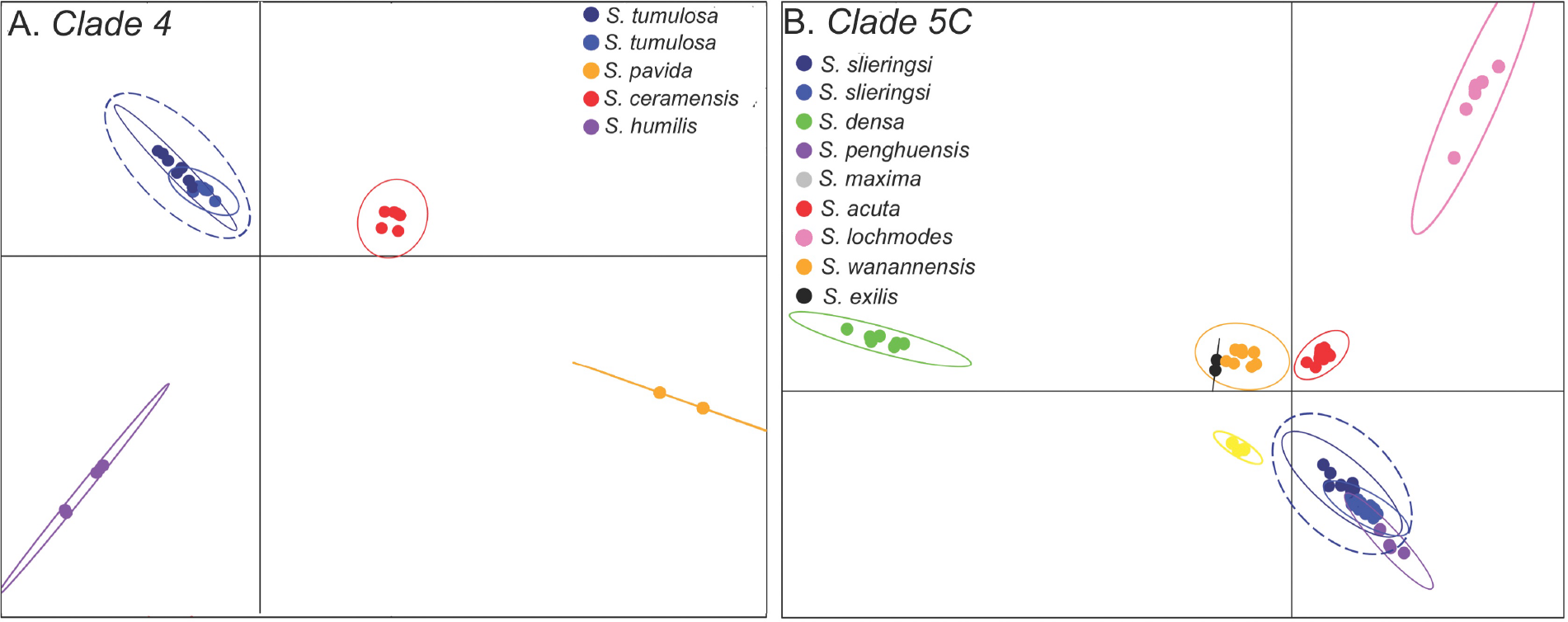
Discriminant analysis of principal components (DAPC) plots for Dongsha Atoll *Sinularia* (A) clade 4 and (B) clade 5C. Genetic clusters representing different morphospecies are color coded to match the phylogenetic trees in Figures 2, 3 and 4. Species (*S. tumulosa* in (A) and *S. slieringsi* in (B) encircled in dotted lines) that were suggested to be further divided into two species by Bayes Factor Determination are also denoted.

Twenty-four separate ABBA-BABA tests were performed on clade 4 (Fig. 7, Suppl. File 1). The average number of loci shared across taxa in each test was 3313 ± 556 (Suppl. File 2). The ABBA-BABA tests indicated admixture between *S. tumulosa* and *S. pavida* lineages (α=3.0, Z= 3.44-4.10, D=0.11; tests 10, 18). Eleven of thirteen individuals of *S. tumulosa* (both clades a and b) appeared to be strongly admixed with *S. pavida* (α=3.0, Z=3.11-4.55, D=0.11-0.18; tests 11, 14-17, 19-24, Suppl. File 2). Upon further examination with partitioned D-statistics, introgression appeared to have occurred from *S. pavida* into both *S. tumulosa* clades (α=3.0, Z = 2.8-3.0, D=-0.14-0.15, Suppl. File 2).

**Figure 7.**
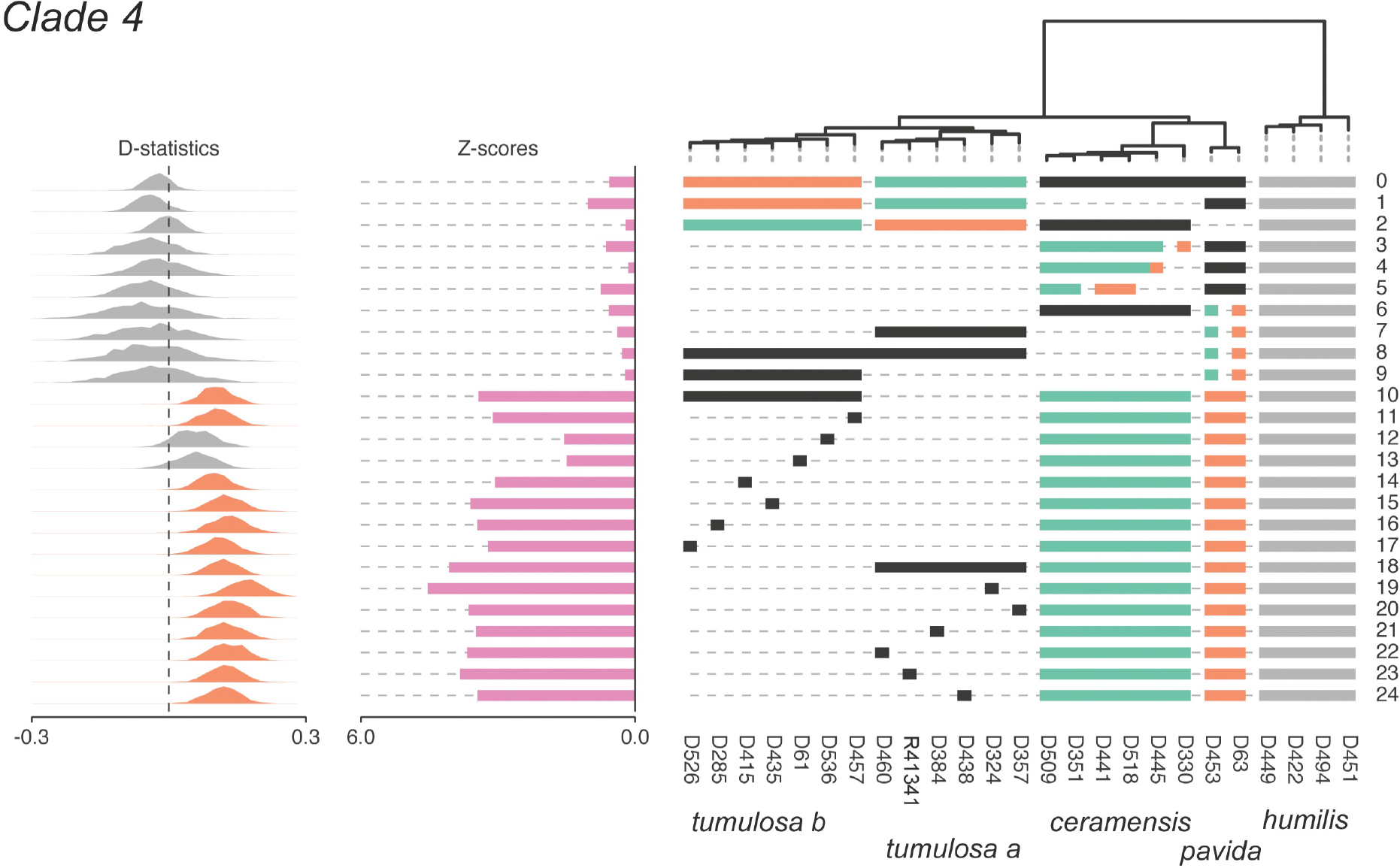
D-statistic tests for admixture in Dongsha Atoll *Sinularia* clade 4. Test numbers are listed on the right for each 4-taxon test (((p1, p2), p3), p4). Horizontal bars below the tips of the tree indicate which taxa were included in each test. *S. humilis* was set as the outgroup for all tests (indicated by gray bars). Tests are configured to ask whether P3 (black bars) shares more derived SNPs with lineage P1 (green bars) relative to P2 (orange bars). As illustrated to the left, Z scores are bar plots and D-statistics are histograms. Histograms are green for significant gene flow between P1 and P3 (BABA) and orange for significant gene flow between P2 and P3 (ABBA). D-statistics that were not significant are gray. Significance was assessed at an alpha level of 3.0 (i.e., when D deviates > 3.0 standard deviations from zero).

### Clade 5C

Species delimitation analyses supported eight species in *Sinularia* clade 5C (*S. acuta/abrupta*, *S. densa*, *S. exilis*, *S. lochmodes*, *S. maxima*, *S. penghuensis/daii*, *S. slieringsi*, and *S. wanannensis*); *S. abrupta* was not distinguished from *S. acuta*, and the holotype of *S. daii* was placed with *S. penghuensis*. In addition, five individuals identified as *S. penghuensis*, including the holotype and paratypes, grouped with *S. slieringsi*. The optimal number of *K* clusters suggested by the DAPC analyses was eight (BIC=377, Suppl. Fig. 2), and the DAPC plot revealed no overlap among these distinct clusters (Fig. 6b). Further support for group assignment can be seen in the assignment plots, as all individuals were successfully re-assigned into their respective clusters (Suppl. Fig. 4). In addition, the Distruct plot clearly illustrated little to no admixture between these eight species (Fig. 4), except that both individuals of *S. exilis* appeared to be admixed with at least three different species, including *S. densa, S. maxima*, and *S. wanannensis*. Individuals of *S. abrupta* (D329) and the holotype of *S. wanannensis* (Z34695) whose *28S* barcode sequences were incongruent with their *mtMutS* haplotypes (Fig. 2b) also showed some evidence of admixture with *S. maxima* and *S. densa*, respectively.

It is possible that *S. slieringsi* represents two cryptic species, although results are not conclusive. Upon further Structure analysis, little to no admixture was revealed between two sub-clades of *S. slieringsi*; however, two individuals (one of them a paratype of *S. penghuensis*, Z34681) were admixed (Fig. 4). Following a one-species model (MLE=−1298), DAPC+1 was the second most likely (MLE=−1543) species model according to BFD* analyses (Table 3). The DAPC+1 model included species designated by DAPC, plus the two groups of *S. slieringsi*. Second, individuals of *S. slieringsi* formed two separate groupings in the DAPC plot, although there was some overlap among individuals (Fig. 6b). *S. slieringsi* also split into two reciprocally monophyletic clades in the species tree phylogeny (Fig. 5b), but not in the concatenated RAxML phylogeny (Fig. 4).

Fifty separate ABBA-BABA tests were run on clade 5C (Fig. 8, Suppl. File 1). The average number of loci shared across taxa in each test was 1745 ± 216 (Suppl. File 2). The ABBA-BABA tests indicated admixture between the *S. penghuensis/S. daii clade* and one clade, clade b, of *S. slieringsi* (α=3.0, Z scores =4.51, D=−0.15; test 14). Most individuals in the latter clade (which included the holotype of *S. penghuensis*) appeared to be strongly admixed as evidenced by ABBA-BABA tests (α=3.0, Z =3.1-5.5, D=−0.13−^-^0.20; tests 15, 19-26, 28). As suggested by the Structure analysis, the holotype of *S. wanannensis* (Z34695) showed strong admixture with *S. densa* (α=3.0, Z=3.13, D=0.14; test 9), but not with *S. maxima* (α=3.0, Z=0.25, D=−0.01; test 2). It was also not admixed with either *S. penghuensis* or *S. slieringsi* (α=3.0, Z=0.24-0.61, D=−0.0.3--0.01; tests 46-47), even though it shared the same *mtMutS* haplotype as both of those species. Although *S. abrupta* D329 showed some evidence of admixture with *S. maxima* and *S. densa* in the Structure analysis, the ABBA-BABA tests indicated that it was not a hybrid (α=3.0, Z=1.59-2.78, D=0.10-0.18; tests 32, 45). The *S. abrupta* specimen Z33623 that shared a *28S* sequence with *S. densa* was also not significantly admixed with that species (α=3.0, Z=2.11, D=0.13, test 44), whereas *S. abrupta* D19 was (α=3.0, Z=312, D=0.15; test 40). One individual of *S. acuta*, D450, was also admixed with *S. densa* (α=3.0, Z=3.21, D=0.16; test 37). Notably, the Distruct plot showed strong admixture of the two *S. exilis* specimens with *S. densa, S. maxima*, and *S. wanannensis* (Fig. 4). However, ABBA-BABA tests suggested that these individuals were not significantly introgressed with those or any other species (α=3.0, Z=0.25-2.96, D=-0.15-0.04; tests 0-2, 5-6, 8, 48-49). Overall, the ABBA-BABA tests indicated that at least 14 individuals were significantly introgressed with other species at an α=3.0; however, we note that D statistics for several other tests also deviated considerably from 0, although they were not significant at α=3.0.

**Figure 8.**
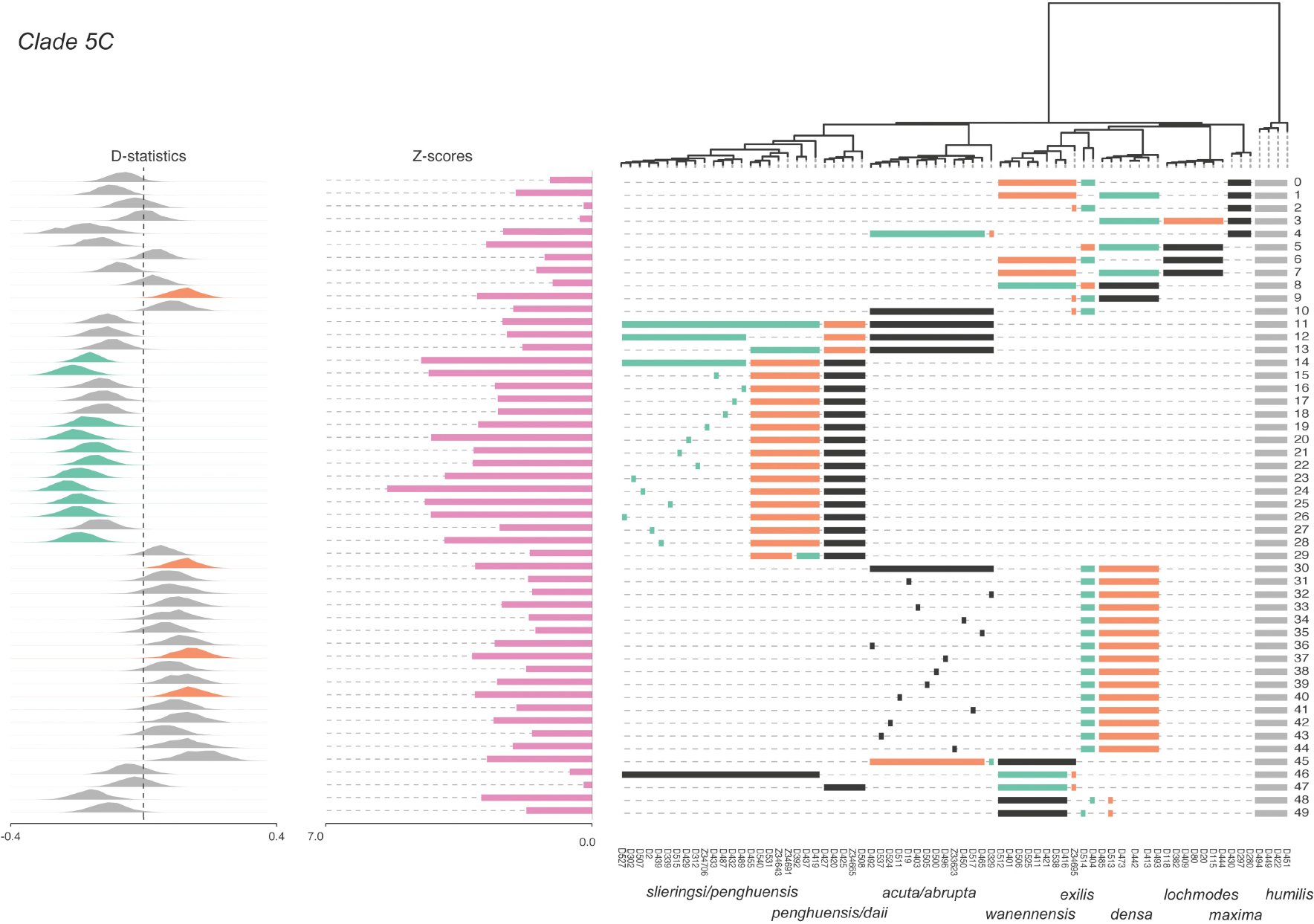
D-statistic tests for admixture in Dongsha Atoll *Sinularia* clade 5C. Test numbers are listed on the right for each 4-taxon test (((p1, p2), p3), p4). Horizontal bars below the tips of the tree indicate which taxa were included in each test. *S. humilis* was set as the outgroup for all tests (indicated by gray bars). Tests are configured to ask whether P3 (black bars) shares more derived SNPs with lineage P1 (green bars) relative to P2 (orange bars). As illustrated to the left, Z scores are bar plots and D-statistics are histograms. Histograms are green for significant gene flow between P1 and P3 (BABA) and orange for significant gene flow between P2 and P3 (ABBA). D-statistics that were not significant are gray. Significance was assessed at an alpha level of 3.0 (i.e., when D deviates > 3.0 standard deviations from zero).

## Discussion

### Species Delimitations

The different species delimitation methods based on RADseq data resulted in incongruence in the number of species suggested to be present among those sampled at Dongsha Atoll in each of *Sinularia* clades 4 and 5C. BFD* analysis indicated that for each clade the most likely model was a one-species model, whereas DAPC indicated that four species were present in clade 4 and eight were present in clade 5C. It seems unlikely that only a single species exists in each of clades 4 and clade 5C because there is little overlap among groups in the DAPC plots and there is strong genetic structure shown in the Structure analyses. Furthermore, there are well-supported clades in both ML and species trees, many of them congruent with distinct morphologies (Benayahu et al., 2018). The suggestion by BFD* analysis of only one species in each of the two clades is likely spurious due to the relatively few loci included in those analyses. Because of the computational time it took to run each SNAPP species-delimitation model, only complete datasets were used, i.e., those that included no missing data, and these contained relatively few SNPs (< 200).

All other species delimitation analyses supported four or five species of *Sinularia* in clade 4 and eight or nine in clade 5C among the samples that were sequenced. Within clade 4, RADseq and both barcoding markers discriminated *S. humilis*, *S. ceramensis* and *S. pavida* from one another and from all specimens identified as *S. tumulosa*. Both the *mtMutS* barcode and RADseq results further delineated two distinct clades within *S. tumulosa*, with hybridization tests suggesting that both are admixed with *S. pavida*. Within clade 5C, species delimitation analyses clearly distinguished *S. lochmodes*, *S. densa*, *S. wanannensis* and *S. maxima* from all other species, and each of those morphospecies also had unique haplotypes at both barcoding loci. *S. acuta* was similarly delineated from all other morphospecies with the notable exception of *S. abrupta*. Two of three individuals of *S. abrupta* shared a *mtMutS* haplotype with *S. acuta*, and could not be distinguished from that species in the DAPC and Structure analyses. A third colony identified as *S. abrupta* (D329) showed signs of admixture in both Structure and ABBA-BABA analyses, suggesting a possible hybrid origin. As *S. acuta* and *S. abrupta* differ markedly in both colony growth form and sclerite morphology (Benayahu et al., 2018), the apparent lack of genetic distinction between these two morphospecies warrants further study.

The remaining morphospecies in clade 5C could not be separated clearly using either barcode marker, and the multilocus analyses suggested that admixture may contribute to the difficulty distinguishing them. Although both individuals of *S. exilis* had unique *mtMutS* haplotypes, they shared 28S genotypes with *S. densa* and S*. wanannensis*. Phylogenetic analyses placed *S. exilis* as the sister to *S. wanannensis*, but Structure suggested considerable admixture with both *S. maxima* and *S. densa*. As ABBA-BABA tests did not strongly support a hybrid origin of *S. exilis*, incomplete lineage sorting may better explain why this species shares genotypes with other species in the clade. Hybridization was, however, supported as a possible explanation for the confusing relationship between *S. penghuensis* and *S. slieringsi*, two morphospecies that shared several different *mtMutS* haplotypes, one of which was also shared by *S. daii*. Five colonies of *S. penghuensis*, including the holotype (ZMTAU Co34706) and two paratypes (ZMTAU Co3464, Co34681), shared a *28S* genotype with *S. slieringsi*, while the other four shared a very different 28S genotype with the holotype of *S. daii* (ZMTAU Co 34665). Multilocus species delimitation analyses separated the latter four *S. penghuensis* plus *S. daii* from a large clade that included all *S. slieringsi* plus the *S. penghuensis* type specimens. Structure and DAPC further separated that large clade into two sub-clades, suggestive of possible cryptic species. ABBA-BABA tests indicated that a majority of the individuals of *S. slieringsi* and *S. penghuensis* in one of those two sub-clades are admixed with the *S. penghuensis-S. daii* clade.

### Evidence for Hybridization in Sinularia

It is important to use a phylogenetic framework in assessments of introgressive hybridization. A species that appears admixed could have a close relative harboring a stronger signal of admixture (Eaton et al., 2015), and if that species is not included in analyses, then the admixture will be incorrectly attributed to a closely-related taxon (Durand et al., 2011; Eaton & Ree, 2013; Rogers & Bohlender, 2015). In addition, it can be challenging to distinguish introgression between two species from “secondary genomic admixture”, which occurs when one species shares recent ancestry with a true hybridizing lineage, thus causing that species to also appear as if it were admixed (Eaton & Ree, 2013; Eaton et al., 2015). Although the current study sequenced most of the clade 4 and 5C morphospecies that occur at Dongsha Atoll, there were at least two morphospecies in each clade that were not included in the analyses. In addition, there are other *Sinularia* species in phylogenetically distinct clades that also inhabit this atoll, and these too were not included in the analyses. While our results provide evidence for hybridization, we acknowledge the possibility of incorrectly attributing admixture to a close relative of the true hybridizing lineage, as not all possible *Sinularia* morphospecies were included in these analyses.

Because of the difficulties interpreting ABBA-BABA tests even when using a phylogenetic framework (Eaton et al., 2015), it is best to focus on the strongest signals of admixture (Eaton, 2018). The signal of introgression was strong in clade 4, with both clades of *S. tumulosa* showing admixture with *S. pavida*; no other hybridization tests in this clade were significant, with D-statistics centered around zero and Z scores fairly low. In contrast, it was a bit more difficult to confidently determine which *Sinularia* species are hybridizing with others in clade 5C, and perhaps this is due to the use of an incomplete phylogeny. There were some cases where the signal of admixture was strong (e.g., *S. acuta* with *S. densa, S. slieringsi* with *S. penghuensis)* and supported by other tests in addition to ABBA-BABA. However, there were also cases where species appeared admixed, but the ABBA-BABA results were not significant at an alpha of 3.0. For example, Structure analyses suggested there was considerable admixture between *S. wanannensis* and *S. exilis*, and at least one *S. exilis* individual had the same *28S* barcode as *S. wanannensis*, but the ABBA-BABA tests were not significant. Perhaps with more *S. exilis* individuals in the analyses, or with the addition of the missing morphospecies of *Sinularia*, a more complete picture would emerge of whether these species share genes as a result of incomplete lineage sorting or hybridization. Because the current phylogenetic analysis did not include all species, it is possible that the species identified as the source of introgressed alleles may simply be close relatives of the actual parental species. Nevertheless, the phylogenetic framework was a useful approach in determining that hybridization appears to be an important process contributing to the diversification of this speciose group of soft corals.

Incongruence between the RAxML and SNAPP species trees (both built using 25% missing data) may provide further support for hybridization among *Sinularia* species. In the clade 4 SNAPP species tree, *S. ceramensis* was sister to a clade of *S. pavida* plus *S. tumulosa*, whereas in the ML tree built using concatenated RAD loci, *S. ceramensis* was sister to *S. pavida*. In clade 5C, *S. acuta* was sister to *S. penghuensis* and *S. slieringsi* in the ML tree, but sister to all other species in the SNAPP species tree. Notably, relationships that differed between analyses showed evidence of admixture in the ABBA-BABA results. Although incomplete lineage sorting can lead to discordance between phylogenies built using concatenated data vs. species tree methods (Edwards et al., 2007; Kubatko & Degnan, 2007; Maddison, 1997), hybridization has also been shown to produce incongruence among gene trees (Edwards et al., 2007; Kubatko, 2009). Introgressive hybridization may also explain the alterative topologies recovered in the SNAPP species tree. Johnston et al. (2017) suggested that alternative trees emerging from SNAPP analyses of corals in the genus *Porites* could be due to introgressive hybridization, incomplete lineage sorting, or contamination by loci of symbionts (e.g., Symbiodiniaceae). Our results, however, indicate that introgressive hybridization likely explains the discordance observed, as the species displaying different relationships in the SNAPP trees were suggested to be hybridizing. Results further lend support to the idea that diversification of species-rich lineages may not be a solely bifurcating process. As such, phylogenetic tree reconstructions that include taxa that do not follow the usual assumption of a bifurcating process of evolution can lead to incongruence among gene trees and contribute to difficulties in resolving phylogenies.

### Morphospecies versus Genetic Data

Incongruence between morphological and molecular evidence for species boundaries is common in corals (e.g., Forsman et al., 2009, 2010; Keshavmurthy et al., 2013). Contributing factors include environmental plasticity (e.g., Paz-García et al., 2015) and frequent homoplasy of morphological characters (e.g., Forsman et al., 2009) as well as the slow rate of mitochondrial gene evolution that has made ‘universal’ molecular barcodes such as COI relatively invariant among congeneric species (Huang et al., 2008; McFadden et al., 2011; Shearer & Coffroth, 2008). When barcodes fail to discriminate distinct morphospecies it may be because the markers lack appropriate variation, or, alternatively, because morphological variation within a species has been incorrectly interpreted as evidence of a species boundary (McFadden et al., 2017). Attempts to integrate the two different sources of evidence have met with some success, as demonstrated by Benayahu et al. (2018). By combining assessment of morphology with a character-based barcoding approach, that study identified at least 27 species of *Sinularia* from Dongsha Atoll, including those used in the current study. In several cases in which distinct morphotypes shared identical *mtMutS* haplotypes, however, they attributed the lack of congruence to invariance of the barcode marker (i.e., incomplete lineage sorting), and delimited species using morphological characters. In two such cases, species delimitation methods using RADseq data also failed to support the genetic distinction between discrete morphotypes, namely *S. acuta* and *S. abrupta*, and *S. slieringsi* and *S. penghuensis*. Moreover, our inclusion of type specimens of *S. penghuensis* and *S. daii* as taxonomic references revealed no genetic distinction between the material identified here as *S. slieringsi* and the *S. penghuensis* types, or between the holotype of *S. daii* and additional colonies identified as *S. penghuensis*. In addition, a colony from Palau identified as *S. verruca* (R41341) could not be distinguished genetically from one of the two clades of *S. tumulosa*. Clearly, additional taxonomic work integrating both morphological and molecular approaches will be necessary to clarify the relationships among these taxa.

Morphological discrimination of species is complicated in *Sinularia* and many other soft corals due to the continuous nature of many of the characters used to diagnose species. Colony growth forms and the intricate shapes of sclerites are difficult to quantify and may present a continuum of variation, as do morphometric characters such as the sizes of sclerites commonly used in the literature (e.g. Verseveldt, 1980). Many of the species examined here, including *S. tumulosa*, *S. verruca, S. acuta* and *S. daii*, were described from single exemplars (Benayahu & Ofwegen, 2011; Manuputty & Ofwegen, 2007; Ofwegen, 2008), and therefore no data exist on the possible range or limits of morphological variation they exhibit, potentially confounding efforts to discriminate them from other similar species. Hybridization also offers a possible explanation for the lack of congruence between morphological and molecular determinations of species identity. As has been suggested for some coral genera, hybridization can lead not only to morphologically distinct or intermediate phenotypes (*Porites*: Forsman et al., 2017; *Acropora*: van Oppen et al., 2000, 2002; Vollmer & Palumbi, 2002), but also to F1 hybrids that exhibit characters of both parental species (Slattery et al., 2008; Vollmer & Palumbi, 2002). In a naturally occurring hybrid zone in Guam, for example, F1 hybrids of *S. maxima* and *S. polydactyla* were found to contain a mix of sclerites resembling those of both parental species (Slattery et al., 2008). Perhaps mechanisms such as these add to the confusion in morphospecies identification, which is often pervasive in *Sinularia* and other octocorals. Although it is currently unknown whether or not hybridization generates new species or asexual lines in the genus *Sinularia*, admixture between *Sinularia* species occurs, and perhaps contributes to the range of morphotypes observed in the genus.

### Utility of DNA Barcoding in Corals

As numerous other studies have now cautioned, none of the single-gene molecular barcodes currently used to help guide species identifications in octocorals successfully delimit all species (Baco & Cairns, 2012; Herrera & Shank, 2016; McFadden et al., 2011, 2014; Pante et al., 2015), particularly when genetic distance thresholds are used to decide species boundaries. For example, in *Sinularia* clade 5C only four MOTUs were identified among eight morphospecies using the *28S* marker, whereas *mtMutS* resolved eight MOTUs, but not all of them were congruent with morphospecies and RADSeq delimitations. Such lack of concordance among different molecular markers is not uncommon in corals and in recently diverged clades more generally (McFadden et al., 2014; Prada et al. 2014; Radice et al., 2016; van Oppen et al. 2001), a result that is likely due to incomplete lineage sorting. For octocorals, a consensus has emerged that mitochondrial and rDNA barcodes may be useful in species assessments for some taxa (e.g., McFadden et al., 2014; Pante et al., 2015), but not all, and that multiple markers and other lines of evidence need to be considered when delimiting species (McFadden et al., 2017). Genomic approaches such as RADseq are effective (Herrera & Shank, 2016; Pante et al., 2015), but still prohibitively expensive and impractical to use for the routine species identification work required of biodiversity surveys. Alternatively, once species boundaries have been validated using such approaches, it may be possible to identify morphological or simple molecular characters that are species-diagnostic. As discussed above, however, the continuum of variation in morphological traits of corals complicates the search for diagnostic characters, and in some recently discriminated octocoral taxa none have yet been identified (McFadden et al., 2017).

Single-gene barcode markers such as *mtMutS* and *28S* offer diagnostic nucleotide characters that can be used to identify cryptic taxa (McFadden et al., 2011). When Benayahu et al. (2018) applied a character-based *mtMutS* barcode to the *Sinularia* species found at Dongsha, the only morphospecies that could not be discriminated were the same ones for which the current study also found incongruence between RAD clades and morphospecies designations: *S. acuta*, *S. abrupta*, *S. penghuensis*, *S. daii* and *S. slieringsi*. A compound, character-based barcode that combines *mtMutS* with *28S*, however, yields diagnostic characters that discriminate each of the *Sinularia* clades identified by RADseq, including both sub-clades of *S. tumulosa* and *S. slieringsi* (Suppl. Fig. 5). Once species boundaries have been validated using integrated, genomic approaches such as those applied here, use of simple character-based barcodes to identify morphologically cryptic species may be more time- and cost-effective than genomic approaches. Assignment of character-based barcodes, however, requires *a priori* recognition of species boundaries, as well as screening of a sufficient number of individuals to identify polymorphic characters.

### Future Research Directions

Further investigation is needed to determine the evolutionary processes responsible for generating the high species diversity in the genus *Sinularia*, but a necessary first step is to understand how many species there are and where they are distributed. With accurate species identifications utilizing both classical taxonomy and advanced genomic techniques, it will be possible to address questions pertaining to how and when *Sinularia* diversified into coral reef environments and why species in this genus appear to be so successful at co-existing on one reef. One example of an intriguing question is whether the high diversity of *Sinularia* was generated in sympatry through mechanisms such as hybrid speciation or whether species have diverged in allopatry and then colonized the same reefs. With the advent of new genomic techniques such as RADSeq and target-capture genomics (e.g., Quattrini et al., 2018), we can begin to examine how pervasive hybridization is on coral reefs, particularly because F1 hybrids and their progeny may be more fit than the parent populations, and hybrid vigor may help in the maintenance and resilience of coral reef diversity (Slattery et al., 2008). Using a genomic approach can help to guide species delimitation while simultaneously shedding light on the processes generating diversity in this genus, just one of many hyperdiverse coral lineages (e.g., the scleractinians *Acropora, Porites, Pocillopora* and the octocorals *Dendronephthya, Lobophytum* and *Sarcophyton*) in which ecologically similar congeners co-occur in high numbers on coral reefs throughout the Indo-Pacific.

## Conclusions

Delimiting species is a critical first step in documenting the biodiversity of ecologically important corals, such as species in the genus *Sinularia*. This study demonstrates the utility of using genomic approaches to delimit species within a hyperdiverse lineage of soft corals and to examine simultaneously whether hybridization may be contributing to its diversification. Although there was some incongruence among datasets and species delimitation methods, we can confidently conclude that the sequenced individuals represent at least four species of *Sinularia* in clade 4 and eight species in clade 5C (McFadden et al. 2009), with the potential for one additional cryptic species in each clade. The results point to hybridization as an important source of diversification in *Sinularia*, and suggest that this mechanism may produce hybrids with morphologies intermediate to those of their parental species, contributing to the difficulty of assigning species based on morphology in this and other coral genera (Forsman et al., 2017; Fukami et al., 2004; Miller & van Oppen, 2003; Slattery et al., 2008). Furthermore, our results raise the possibility that hybrid speciation (i.e., reticulate evolution via introgressive hybridization) is one mechanism that has contributed to the diversity of octocorals.

## Methods

### Specimen Collection and Preparation

Colonies of *Sinularia* were collected using SCUBA during biodiversity surveys conducted at Dongsha Atoll Marine National Park (Taiwan) in 2011, 2013 and 2015 (Benayahu et al., 2018). Collections were made at 13 sites in the shallow fore-reef zone (3-21 m) surrounding the 25-km diameter atoll (Fig. 1). During the 2015 survey we specifically targeted common morphospecies belonging to clades 4 and 5C (McFadden et al., 2009). Following collection, small subsamples of tissue were preserved in 95% EtOH, and the remainder of the specimen was preserved in 70% EtOH. All vouchers have been deposited in the Steinhardt Museum of Natural History, Tel Aviv University, Israel (ZMTAU, Suppl. File 3). To identify morphospecies, sclerites were isolated from colonies by dissolving tissue in 10% sodium hypochlorite and examined using either light microscopy or, when necessary, scanning electron microscopy (SEM) (Benayahu et al., 2018). The morphological IDs were made by direct comparison to type material when available. Specimens belonging to 13 *Sinularia* morphospecies, four from clade 4 and nine from clade 5C, were selected for further species delimitation analyses. Seven specimens collected in previous biodiversity surveys of the Penghu Archipelago, Taiwan (Benayahu et al., 2012) and Palau (McFadden et al., 2014), including original type material of *S. penghuensis* (ZMTAU Co34643, Co34706, Co34681), *S. wanannensis* (ZMTAU Co34695) and *S. daii* (ZMTAU Co34665), were also included as taxonomic references (Table 1; Suppl. File 3).

DNA was extracted from 95 *Sinularia* specimens, and quantified using a Qubit v 2.0 fluorometer (Broad Range Assay Kit). Quality was assessed by running 100 ng of DNA for each sample on a 1% agarose gel, and checked with a Nanodrop spectrophotometer. Concentration of high-quality (230/260 and 260/280 ratios >1.8) DNA was normalized to 20 ng per ul and sent to Floragenex Inc (Eugene, OR) for RADSeq library preparation. DNA libraries were constructed for each of the 95 samples using the 6-cutter *PstI* enzyme, and then split into two for sequencing 100 bp SE reads on two full lanes of an Illumina HiSeq2500 (University of Oregon’s Genomics and Cell Characterization Core Facility lab). In addition to RADseq, two gene regions (*mtMutS*, *28S* rDNA) used widely for barcoding in octocorals were PCR-amplified and Sanger-sequenced using published primers and protocols (McFadden et al., 2011, 2014).

### Phylogenetic Inference and Species Delimitation using DNA Barcodes

*mtMutS* and 28S sequences of the *Sinularia* species were each aligned using the L-INS-i method in MAFFT (Katoh & Toh, 2008), and pairwise genetic distances (Kimura 2-parameter) among sequences were calculated using the DNADist program in PHYLIP v. 3.69 (Felsenstein, 2005). MOTHUR v 1.29 (Schloss et al., 2009) was used to delimit molecular operational taxonomic units (MOTUs) based on a genetic distance threshold of 0.3% applied separately to *mtMutS* and 28S (e.g., McFadden et al., 2014). Phylogenetic trees were constructed separately for *mtMutS* and *28S* rDNA using maximum likelihood methods (Garli; Zwickl, 2006) (Suppl. Fig. 1). jModelTest (Darriba et al., 2012) was used to identify the best models of evolution (AIC criterion) to use in these analyses (*mtMutS*: HKY + G; *28S rDNA*: HKY + I).

### Phylogenetic Inference and Species Delimitation using RADSeq Data

We produced several different RADSeq locus datasets from the *Sinularia* material (Suppl. Files 4-5; all tree files on datadryad). Datasets chosen for species delimitation and phylogenetic analyses included those loci from pyRAD parameter settings that maximized the number of phylogenetically informative sites and reduced the chances for including paralogous loci (Table 2) Thus, for most analyses, we used data produced from the following parameter settings: *c* 0.85, *p* 0.25, and *m* 0.75 for each of clades 4 and 5C.

RAxML v8 (Stamatakis, 2006) was used to create maximum likelihood (ML) phylogenies for clades 4 and 5C. A GTR+G+I model as suggested by the Akaike Information Criterion [(AIC=371918), JModelTest v2, Darriba et al., 2012] was used. A total of 20 ML searches and 200 bootstrap replicates were performed using rapid bootstrapping on concatenated loci. RAxML analyses were performed (12 analyses per clade) for each of the different datasets produced by pyRAD with different parameter combinations and clustering thresholds (Table 2, Suppl. Files 4-5).

Discriminant Analysis of Principal Components (DAPC) was performed using the package ‘adegenet’ v2.0 in R (Jombart & Collins, 2015; R Core Team, 2012) to explore genetic structure in both clades 4 and 5C. The DAPC method, used previously in species delimitation analyses (Pante et al., 2015), forms clusters based on genetic similarity of each multilocus genotype, without considering a model of evolution. We first used the function *find.clusters* to find the best number of *K* genetic clusters in unlinked SNP datasets for each clade. *Find.clusters* was performed on *K*=20 for each clade. The lowest value of the Bayesian Information Criterion (BIC) statistic was used to detect the optimal number of *K* clusters. These clusters were then analyzed using DAPC, which first transforms the data using principal components analysis and then performs a Discriminant Analysis on the retained principal components. The *optim.a.score* function was used to determine how many PC axes needed to be retained (Suppl. Fig. 6). Six PCAs and four discriminant functions were retained for clade 4; and six PCAs and six discriminant functions were retained for clade 5C (Suppl. Fig. 7). Scatterplots of discriminant functions were then created. We also used the function *assignplot* to visualize individual membership in each *K* cluster, which can help show the accuracy of the cluster assignments and identify any individuals that have high probabilities of membership in >1 cluster.

The Bayesian model-based clustering approach, Structure v2.3 (Pritchard et al., 2000), was also used to infer the number of *Sinularia* species. The program clusters individuals based on genetic variation alone, without any other prior information such as geographic origin or population assignment. Structure was performed on unlinked SNP datasets for both *Sinularia* clades 4 and 5C, and run in parallel using StrAutoParallel v 1.0 (Chhatre & Emerson, 2017) using an admixture model with correlated allele frequencies. Burnin was set to 250,000 followed by 1,000,000 MCMC generations. The inferred number of populations (*K*) was set from 1 to 8 for clade 4 and 1 to 10 for clade 5C; 20 runs of each *K* were conducted. We plotted the population structure assignments of *K=4* for clade 4 and *K=8* for clade 5C, because these were the number of genetic clusters suggested by DAPC analyses. Multiple runs of K were aligned with CLUMPP v 1.2 (Jakobsson & Rosenberg, 2007), and the resulting *indivq* file was input into Distruct v. 1.1 (Rosenberg, 2004) for graphical display of individual population assignments. The commonly used ∆K method (Evanno et al., 2005) was not used in our study to identify an optimal *K* value because this method is known to be most successful at finding only the uppermost levels of genetic structure in a hierarchical system (Evanno et al., 2005). In addition, initial tests of the ∆*K* method revealed *K*=2 as the best model for each clade; however, analysis at *K*> 2 indicated strong genetic structure in both clades 4 and 5C. Therefore, following Gowen et al., (2014), we analyzed successively smaller groups of species in separate analyses. We re-ran structure on the putative species, *S. tumulosa* and *S. slieringsi*, because Bayes Factor Delimitation with genome data (BFD*) analyses (see below), suggested the presence of additional species within each of *S. tumulosa* and *S. slieringsi*.

Coalescent-based SNAPP v 1.3 (Bryant et al., 2012) analyses were used to test alternative species models for both clades 4 and 5C. Samples were assigned to the following alternative species models (Suppl. Figs. 8-9): 1) one species (ONESP), 2) two species (TWOSPP), 3) DAPC clusters (DAPC), 4) DAPC clusters plus division of another clade (DAPC+1), 5) MOTHUR species assignments based on *mtMutS* (MUTS), and 6) MOTHUR species assignments based on *28S* rDNA (28S). In addition, three (THREESPP) and four (FOURSPP) species models were also tested for clades 4 and 5C, respectively. SNAPP analyses were performed in BEAST v 2.4.5 (Bouckaert et al., 2014) with a path sampling of 48 steps (MCMC length=100,000, pre-burnin= 1,000) following Leaché et al. (2014) and Herrera & Shank (2016). The *c*0.85, *m*1.0, *p*0.25 bi-allelic SNP datasets (175 SNPs for clade 4, 140 SNPs for clade 5C, no missing data) were used because of the long computational time it took for each SNAPP run. Marginal likelihood estimates were obtained for each different model run in SNAPP analyses. The different species delimitation models were then ranked using BFD* methods. Bayes Factors were calculated between each of two alternative models by subtracting the marginal likelihood estimates between two models, and then multiplying the difference by two (following Kass & Raftery, 1995; Leaché et al., 2014).

SNAPP was also used to infer the species tree for each *Sinularia* clade. Three independent runs were performed on SNP data (MCMC length=1,000,000, pre-burnin=1,000, samplefreq=1,000) using BEAST with default parameters for mutation rate, coalescent rate, and ancestral population sizes (following Herrera & Shank, 2016). The *c*0.85, *m*0.75, *p*0.25 bi-allelic SNP datasets were used for species tree analyses. Acceptance probabilities were checked to ensure that tuning parameters were appropriate and the chain mixed well (Drummond & Bouckaert, 2015). Log files were combined using LogCombiner v 1.1 and input into Tracer v1.6 (Rambaut & Drummond, 2007). Convergence and ESS>200 were assessed using Tracer after a 10% burnin. Maximum clade credibility trees were generated with TreeAnnotator v 2.3 (Bouckaert et al., 2014). Both the consensus tree and all tree topologies were drawn in DensiTree v2.2 (Bouckart, 2010).

### Hybridization Tests

We calculated Patterson’s D statistics in ipyRAD v 0.7.28 (Eaton & Overcast, 2016) to test for hybridization between species. Briefly, these tests calculate the proportion of ABBA and BABA site patterns, and excess of either is indicative of admixture rather than incomplete lineage sorting (Durand et al., 2011; Green et al., 2010). Multiple 4-taxon tests were generated for both clades 4 and 5C (Suppl. File 1). For both clades, *S. humilis* was set as the outgroup (‘p4’). For tests that included multiple individuals per lineage, SNP frequencies were pooled. For tests performed on clade 4, each species was set as ‘p3’ and all possible 4-species combinations were tested. For clade 5C, all possible 4-species combinations were tested in each of two sub-clades (i.e., the *S*. *slieringsi-S.penghuensis*-*S. acuta* clade and the *S*. *wanannensis-S.exilis-S. lochmodes-S. densa*-S. *maxima* clade). Additional tests were conducted either when Structure results indicated potential admixture or there was incongruence between the different molecular markers (i.e., 28S, *mtMutS*, RAD). When test results were significant at the species level, further tests were performed to determine if particular individuals within the lineage were admixed. Significance of each test was determined by performing 1000 bootstrap replicates in which loci were resampled with replacement. Both D statistics and Z statistics, which represent the number of bootstrap standard deviations (alpha=3.0) that D statistics deviate from zero (Federman et al., 2018), are reported. Following D-statistic tests, partitioned D-statistic tests were performed for a few cases to examine the direction of introgression. Tests were conducted and figures were plotted following the ipyRAD ABBA-BABA cookbook in Jupyter Notebook (Kluyver et al., 2016; see Suppl. File 6).

## Abbreviations

RADSeq: Restriction Site-Associated Sequencing

## Declarations

### Ethics Approval

Not Applicable

### Consent for Publication

Not Applicable

## Data Availability

Raw RADSeq data: SRA#####

Mafft alignments: Data Dryad###

.Tre files: Data Dryad ####

28S sequences: Genbank #s MK333539–MK333628

mtMutS: GenBank #s: See Suppl. File 1

## Competing Interests

The authors declare that they have no competing interests.

## Funding

This study was made possible by a grant to YB from the Taiwanese Ministry of Science (MOST) to conduct octocoral surveys at Dongsha Atoll. Additional funding was provided by Dongsha Atoll Research Station and Howard Hughes Medical Institute Undergraduate Science Education Program award #52006301 to Harvey Mudd College.

## Author Contributions

CSM, TW and YB conceived and designed the study. AMQ and CSM conducted all phylogenetic and species delimitation analyses, and co-wrote the manuscript. TW prepared samples, barcoded genes and conducted preliminary RADSeq analyses. YB made morphological identifications. CSM and YB collected specimens, facilitated by KS and MSJ. All authors edited and approved the final version of this manuscript.

## Acknowledgements

All collections were made in accordance with permits obtained from Marine National Parks, Taiwan. We thank the staff members of Dongsha Atoll National Park, Dongsha Atoll Research Station (DARS), and Biodiversity Research Center, Academia Sinica (BRCAS) for assistance during the field work; A. Gonzalez, J.M. Adams, A. Torres-Navarro, and N. Liu for help with DNA extraction and barcoding; Ben Titus for helpful discussions; and Alex Shlagman for curatorial skills.

## References

Aratake S, Tomura T, Saitoh S, Yokokura R. Kawanishi Y., Shinjo R, Reimer JD, Tanaka J, Maekawa H. Soft coral Sarcophyton (Cnidaria: Anthozoa: Octocorallia) species diversity and chemotypes. PLoS One. 2012;7:e30410.

Babcock RC, Bull GD, Harrison PD, Heyward AJ, Oliver JK, Wallace CC, Willis BL. Synchronous spawnings of 105 scleractinian coral species on the Great Barrier Reef. Mar Bio. 1986;90:379–394.

Baco AR, Cairns SD. Comparing molecular variation to morphological species designations in the deep-sea coral Narella reveals new insights into seamount coral ranges. PLoS One. 2012;7: e45555.

Benayahu Y, Loya Y. Space partitioning by stony corals, soft corals and benthic algae on the coral reefs of the northern Gulf of Eilat (Red Sea). Helgolander wissenschaftliche Meeresuntersuchungen. 1977;30:362–382.

Benayahu Y. Species composition of soft corals (Octocorallia, Alcyonacea) on the coral reefs of Sesoko Island, Ryukyu Archipelago, Japan. Galaxea. 1995;12:103–124.

Benayahu Y, Ofwegen LP van. New species of octocorals (Coelenterata: Anthozoa) from the Penghu Archipelago, Taiwan. Zool. Studies, 2011;50:350–362.

Benayahu Y, Ofwegen LP van, Soong K, Dai CF, Jeng MS, Shlagman A, Hsieh HJ, McFadden CS. Diversity and distribution of Octocorals (Coelenterata: Anthozoa) on the coral reefs of Penghu, Taiwan. Zool. Studies. 2012;51:1529–1548.

Benayahu Y, Ofwegen LP van, Dai CF, Jeng MS, Soong K, Shlagman A, …McFadden CS. The octocorals of Dongsha Atoll (South China Sea): an iterative approach to species identification using classical taxonomy and molecular barcodes. Zool. Studies. 2018;57:doi:10.6620/ZS.2018.57-50.

Blunt JW, Carroll AR, Copp BR, Davis RA, Keyzers RA, Prinseps MR. Marine natural products. Natural Products Reports. 2018;35:8–53

Bouckaert RR. DensiTree: making sense of sets of phylogenetic trees. Bioinformatics. 2010;26:1372–1373

Bouckaert R, Heled J, Kühnert D, Vaughan T, Wu CH, Xie D, Drummond AJ. BEAST 2: a software platform for Bayesian evolutionary analysis. PLoS Comp. Biol. 2014;10:4:e1003537

Bruno JF, Siddon CE, Witman JD, Colin PL, Toscano MA. El Niño related coral bleaching in Palau Western Caroline Islands. Coral Reefs. 2001;20:127–136

Bryant D, Bouckaert R, Felsenstein J, Rosenberg NA, RoyChoudhury A. Inferring species trees directly from biallelic genetic markers: bypassing gene trees in a full coalescent analysis. Mol. Biol. Evol. 2012;29:1917–1932

Carlo JM, Barbeitos MS, Lasker HR. Quantifying complex shapes: elliptical Fourier analysis of octocoral sclerites. Biol. Bull. 2011;220:224–237

Catchen J, Hohenlohe PA, Bassham S, Amores A, Cresko WA. Stacks: an analysis tool set for population genomics. Mol. Ecol‥ 2013;22:3124–3140

Chhatre VE, Emerson KJ. StrAuto: automation and parallelization of STRUCTURE analysis. BMC Bioinformatics. 2017;18:192

Combosch DJ, Vollmer SV. Trans-Pacific RAD-Seq population genomics confirms introgressive hybridization in Eastern Pacific Pocillopora corals. Mol. Phy. Evol. 2015;88:154–162

Currat M, Ruedi M, Petit RJ, Excoffier L. The hidden side of invasions: Massive introgression by local genes. Evolution. 2008;62:1908–1920

Darriba D, Taboada GL, Doallo R, Posada D. jModelTest 2: more models new heuristics and parallel computing. Nature Methods. 2012;9:772

Diekmann O, Bak R, Stam W, Olsen J. Molecular genetic evidence for probable reticulate speciation in the coral genus Madracis from a Caribbean fringing reef slope. Mar. Biol. 2001;139:221–233

Dinesen ZD. Patterns in the distribution of soft corals across the Great Barrier Reef. Coral Reefs. 1983;1:229–236

Drummond AJ, Bouckaert RR. Bayesian evolutionary analysis with BEAST. Cambridge University Press

Durand EY, Patterson N, Reich D, Slatkin M. Testing for ancient admixture between closely related populations. Mol. Biol. Evol. 2011;28:2239–2252

Eaton DA. PyRAD: assembly of de novo RADseq loci for phylogenetic analyses. Bioinformatics. 2014;30:1844–1849

Eaton DA, Ree RH. Inferring phylogeny and introgression using RADseq data: an example from flowering plants (Pedicularis: Orobanchaceae). Syst. Biol. 2013;62:689–706

Eaton DA, Hipp AL, González‐Rodríguez A, Cavender‐Bares J. Historical introgression among the American live oaks and the comparative nature of tests for introgression. Evolution. 2015;69:2587–2601

Eaton DAR, Overcast I. ipyRAD: interactive assembly and analysis of RADseq data sets. 2016

Eaton DA. Ipyrad Cookbook. 2018. http://nbviewerjupyterorg/github/dereneaton/ipyrad/blob/master/tests/cookbook-abba-babaipynb. accessed 20 December 2018

Edwards SV, Liu L, Pearl DK. High-resolution species trees without concatenation. PNAS. 2007;104:5936–5941

Evanno G, Regnaut S, Goudet J. Detecting the number of clusters of individuals using the software STRUCTURE: a simulation study. Mol. Ecol. 2005;14:2611–2620

Fabricius KE. Reef invasion by soft corals: which taxa and which habitats? In: Greenwood JG, Hall NJ, editors. Proceedings of the Australian Coral Reef Society 75th Anniversary Conference Heron Island October 1997. University of Queensland Brisbane; 1998. P. 77–90

Fabricius K. Tissue loss and mortality of soft corals following mass-bleaching. Coral Reefs. 1999;18:54

Fabricius K, Alderslade P. Soft corals and sea fans: a comprehensive guide to the tropical shallow-water genera of the Central West-Pacific the Indian Ocean and the Red Sea. Townsville: Australian Institute of Marine Science; 2001

Federman S, Donoghue MJ, Daly DC, Eaton DA. Reconciling species diversity in a tropical plant clade Canarium Burseraceae. PloSOne. 2018;13:6:e0198882

Felsenstein J. PHYLIP phylogeny inference package Distributed by the author. Department of Genome Sciences University of Washington Seattle Version 3; 2005

Forsman ZH, Barshis DJ, Hunter CL, Toonen RJ. Shape-shifting corals: molecular markers show morphology is evolutionarily plastic in Porites. BMC Evol. Biol‥ 2009;9:45

Forsman ZH, Concepcion GT, Haverkort RD, Shaw RW, Maragos JE, Toonen RJ. Ecomorph or endangered coral? DNA and microstructure reveal Hawaiian species complexes: Montipora dilatata/flabellata/turgescens M. patula/verrilli. PLoS One. 2010;5:e15021

Forsman ZH, Knapp ISS, Tisthammer K, Eaton DAR, Belcaid M, Toonen RJ. Coral hybridization or phenotypic variation? Genomic data reveal gene flow between Porites lobata and P. compressa. Mol. Phyl. Evol. 2017;111:132–148

Frade PR, Reyes-Nivia MC, Faria J, Kaandorp JA, Luttikhuizen PC, Bak RPM. Semi-permeable species boundaries in the coral genus Madracis: introgression in a brooding coral system. Mol. Phyl. Evol. 2010;57:1072–1090

Fukami H, Budd AF, Levitan DR, Jara J, Kersanach R, Knowlton N. Geographic differences in species boundaries among members of the Montastraea annularis complex based on molecular and morphological markers. Evolution. 2004;58:324–337

Goulet TL, LaJeunesse TC, Fabricius KE. Symbiont specificity and beaching susceptibility among soft corals in the 1998 Great Barrier Reef mass coral bleaching event. Mar. Biol. 2006;154:795–804

Gowen FC, Maley JM, Cicero C, Peterson AT, Faircloth BC, Warr TC, McCormack JE. Speciation in Western Scrub-Jays Haldane’s rule and genetic clines in secondary contact. BMC Evol. Biol‥ 2014;14:135

Green RE, Krause J, Briggs AW, Maricic T, Stenzel U, Kircher M, Hansen NF. A draft sequence of the Neandertal genome. Science 2010;328:710–722

Harrison PL, Babcock RC, Bull GD, Oliver JK, Wallace CC, Willis BL. Mass spawning in tropical reef corals. Science 1984;223:1186–1189

Hatta M, Fukami H, Wang W, Omori M, Shimoike K, Hayashibara T, Sugiyama T. Reproductive and genetic evidence for a reticulate evolutionary history of mass-spawning corals. Mol. Biol. Evol. 1999;16:1607–1613

Hebert PD, Penton EH, Burns JM, Janzen DH, Hallwachs W. Ten species in one: DNA barcoding reveals cryptic species in the neotropical skipper butterfly Astraptes fulgerator. PNAS. 2004a;101:14812–14817

Hebert PD, Stoeckle MY, Zemlak TS, Francis CM. Identification of birds through DNA barcodes. PLoS Biology. 2004b;2:e312

Herrera S, Shank TM. RAD sequencing enables unprecedented phylogenetic resolution and objective species delimitation in recalcitrant divergent taxa. Mol. Phyl. Evol. 2016;100:70–79

Hickerson MJ, Meyer CP, Moritz C. DNA barcoding will often fail to discover new animal species over broad parameter space. Syst. Biol. 2006;55:729–739

Huang D, Meier R, Todd PA, Chou LM. Slow mitochondrial COI sequence evolution at the base of the metazoan tree and its implications for DNA barcoding. JJ. Mol. Evol. 2008;66:167–174

Jakobsson M, Rosenberg NA. CLUMPP: a cluster matching and permutation program for dealing with label switching and multimodality in analysis of population structure. Bioinformatics. 2007;23:1801–1806

Jeng M-S, Huang H-D. Dai C-F, Hsiao Y-C, Benayahu Y. Sclerite calcification and reef-building in the fleshy octocoral genus Sinularia (Octocorallia: Alcyonacea). Coral Reefs. 2011;30:925–933

Johnston EC, Forsman ZH, Flot JF, Schmidt-Roach S, Pinzón JH, Knapp IS, Toonen RJ. A genomic glance through the fog of plasticity and diversification in Pocillopora. Scientific Reports. 2017;7:5991

Jombart T, Collins C. A tutorial for discriminant analysis of principal components DAPC using adegenet. Imperial College of London MRC Center for Outbreak Analysis and Modelling; 2015

Kahng SE, Benayahu Y, Lasker HR. Sexual reproduction in octocorals. Mar. Ecol. Prog. Ser. 2011;443:265–283

Katoh K, Toh H. Recent developments in the MAFFT multiple sequence alignment program. Briefings in Bioinformatics 2008;9:286–298

Kass RE, Raftery AE. Bayes factors. J. Am. Stat. Assoc. 90. 1995;90:773–795

Keshavmurthy S, Yang S-Y, Alamaru A, Chuang Y-Y, Pichon M, Obura D, Chen AC. DNA barcoding reveals the coral ‘laboratory rat’ Stylophora pistillata encompasses multiple identities. Scientific Reports. 2013;3:1520

Kim E, Lasker HR, Coffroth MA, Kim K. Morphological and genetic variation across reef habitats in a broadcast-spawning octocoral. Hydrobiologia 2004;530:423–432

Kluyver T, Ragan-Kelley B, Pérez F, Granger BE, Bussonnier M, Frederic J, Ivanov P. May Jupyter Notebooks-a publishing format for reproducible computational workflows. In: ELPUB; 2016. p. 87–90

Kubatko LS, Degnan JH. Inconsistency of phylogenetic estimates from concatenated data under coalescence. Syst. Biol. 2007;56:17–24

Kubatko LS. Identifying hybridization events in the presence of coalescence via model selection. Syst. Biol. 2009; 58:478–488

LaJeunesse TC, Parkinson JE, Gabrielson PW, Jeong HJ, Reimer JD, Voolstra CR, Santos SR. Systematic revision of Symbiodiniaceae highlights the antiquity and diversity of coral endosymbionts. Current Biol. 2018;28:2570–2580

Leaché AD, Fujita MK, Minin VN, Bouckaert RR. Species delimitation using genome-wide SNP data. Syst. Biol. 2014;63:534–542

Lin S, Cheng S, Song B, Zhong X, Lin X, Li W, Cai M. The Symbiodinium kawagutii genome illuminates dinoflagellate gene expression and coral symbiosis. Science. 2015;350:691–694

Maddison WP. Gene trees in species trees. Systematic biology 1997;46:523–536

Manuputty AEW, Ofwegen LP van. The genus Sinularia (Octocorallia: Alcyonacea) from Ambon and Seram Moluccas Indonesia Zoologische Mededelingen Leiden. 2007;81:187–216

Marshall PA, Baird AH. Bleaching of corals on the Great Barrier Reef: differential susceptibilities among taxa. Coral Reefs. 2000;19:155–163

McFadden CS, Hutchinson MB. Molecular evidence for the hybrid origin of species in the soft coral genus Alcyonium (Cnidaria: Anthozoa: Octocorallia). Mol. Ecol. 2004;13:1495–1505

McFadden CS, Ofwegen LP van, Beckman EJ, Benayahu Y, Alderslade P. Molecular systematics of the speciose Indo-Pacific soft coral genus Sinularia (Anthozoa: Octocorallia). Invert. Biol. 2009;128:303–323

McFadden CS, Benayahu Y, Pante E, Thoma JN, Nevarez PA, France SC. Limitations of mitochondrial gene barcoding in Octocorallia. Mol. Ecol. Res. 2011;11: 19–31

McFadden CS, Brown AS, Brayton C, Hunt CB, Ofwegen LP van. Application of DNA barcoding to biodiversity studies of shallow-water octocorals: molecular proxies agree with morphological estimates of species richness in Palau. Coral Reefs. 2014;33:275–286

McFadden CS, Haverkort-Yeh R, Reynolds AM, Halàsz A, Quattrini AM, Forsman ZH, Toonen RJ. Species boundaries in the absence of morphological ecological or geographical differentiation in the Red Sea octocoral genus Ovabunda (Alcyonacea: Xeniidae). Mol. Phyl. Evol. 2017;112:174–184

Miller DJ, Van Oppen MJ. A ‘fair go’ for coral hybridization. Mol. Ecol. 2003;12:805–807

Ofwegen LP van. Status of knowledge of the Indo-Pacific soft coral genus Sinularia May 1898 (Anthozoa: Octocorallia) Proceedings of the 9th International Coral Reef Symposium. 2002;1:167–171

Ofwegen LP van. The genus Sinularia (Octocorallia: Alcyonacea) at Palau Micronesia. Zoologische Mededelingen Leiden. 2008;82:631–735

Ofwegen LP van, Benayahu Y, McFadden CS. Sinularia leptoclados Ehrenberg 1834 Cnidaria: Octocorallia re-examined. ZooKeys. 2013;272:29–59

Ofwegen LP van, McFadden CS, Benayahu Y. Sinularia polydactyla Ehrenberg 1834 (Cnidaria, Octocorallia) re-examined with description of a new species. ZooKeys. 2016;581:71–126

Pante E, Abdelkrim J, Viricel A, Gey D, France SC, Boisselier MC, Samadi S. Use of RAD sequencing for delimiting species. Heredity 2015;114:450

Paz-García DA, Hellberg ME, García-de-León FJ, Balart EF. Switch between morphospecies of Pocillopora corals. Am. Nat. 2015;186:434–440

Prada C, DeBiasse MB, Neigel JE, Yednock B, Stake JL, Forsman ZH, Hellberg ME. Genetic species delineation among branching Caribbean Porites corals. Coral Reefs. 2014;33:1019–1030

Pritchard JK, Stephens M, Rosenberg NA, Donnelly P. Association mapping in structured populations. The American Journal of Human Genetics. 2000;67:170–181

Quattrini AM, Faircloth BC, Dueñas LF, Bridge TC, Brugler MR, Calixto‐Botía IF, et al. McFadden CS. Universal target‐enrichment baits for anthozoan (Cnidaria) phylogenomics: New approaches to long‐standing problems. Mol. Ecol. Res. 2018;18:281–295

R Core Team. R: A Language and Environment for Statistical Computing. R Foundation for Statistical Computing Vienna Austria. 2012 http://wwwr-projectorg/, accessed 15 Jan 2019.

Radice VZ, Quattrini AM, Wareham VE, Edinger EN, Cordes EE. Vertical water mass structure in the North Atlantic influences the bathymetric distribution of species in the deep-sea coral genus Paramuricea. Deep Sea Res. I. 2016;116: 253–263

Rambaut A, Drummond AJ. Tracer v1.4 2007; 2012

Richards ZT, van Oppen MJH, Wallace CC, Willis BL, Miller DJ. Some rare Indo-Pacific coral species are probable hybrids. PLoS One. 2008;3:e3240

Richmond RH, Hunter CL. Reproduction and recruitment of corals: comparisons among the Caribbean the Tropical Pacific and the Red Sea. Mar. Ecol. Prog. Ser. 1990;60:185–203

Rheindt FE, Edwards SV. Genetic introgression: an integral but neglected component of speciation in birds. The Auk. 2011;128:620–632

Rogers AR, Bohlender RJ. Bias in estimators of archaic admixture. Theoretical Pop. Biol. 2015;100:63–78

Rosenberg NA. DISTRUCT: a program for the graphical display of population structure. Mol. Ecol. Notes. 2004;4:137–138

Rowley SJ, Pochon X, Watling L. Environmental influences on the Indo–Pacific octocoral Isis hippuris Linnaeus 1758 Alcyonacea: Isididae: genetic fixation or phenotypic plasticity? PeerJ 2015;3:e1128

Sánchez JA, Aguilar C, Dorado D, Manrique N. Phenotypic plasticity and morphological integration in a marine modular invertebrate. BMC Evol. Biol. 2007;7: 122

Schloss PD, Westcott SL, Ryabin T, Hall JR, Hartmann M, Hollister EB, Sahl JW. Introducing mothur: open-source platform-independent community-supported software for describing and comparing microbial communities. Appl. Envir. Micro. 2009;75:7537–7541

Schmieder R, Edwards R. Fast identification and removal of sequence contamination from genomic and metagenomic datasets. PLoS One. 2011;6: e17288

Shearer TL, Coffroth MA. Barcoding corals: limited by interspecific divergence not intraspecific variation. Mol. Ecol. Res. 2008;8:247–255

Shoham E, Benayahu Y. Higher species richness of octocorals in the upper mesophotic zone in Eilat Gulf of Aqaba compared to shallower reef zones. Coral Reefs. 2017;36:71–81

Shoham E, Prohaska T, Barkai Z, Zitek A, Benayahu Y. Soft corals form aragonite-precipitated columnar spiculite in mesophotic reefs. Scientific Reports. In press

Slattery M, Hines GA, Starmer J, Paul VJ. Chemical signals in gametogenesis spawning and larval settlement and defense of the soft coral Sinularia polydactyla. Coral Reefs. 1999;18:75–84

Slattery M, Starmer J, Paul VJ. Temporal and spatial variation in defensive metabolites of the tropical soft corals Sinularia maxima and S. polydactyla. Mar. Biol. 2001;138:1183–1193

Slattery M, Kamel HN, Ankisetty S, Gochfield DJ, Hoover CA, Thacker RW. Hybrid vigor in a tropical Pacific soft-coral community. Ecol Monogr. 2008;78:423–443

Stamatakis A. RAxML-VI-HPC: maximum likelihood-based phylogenetic analyses with thousands of taxa and mixed models. Bioinformatics. 2006;22:2688–2690

Sukumaran J, Knowles LL. Multispecies coalescent delimits structure not species. PNAS. 2017;114:1607–1612

Titus BM, Daly M. Reduced representation sequencing for symbiotic anthozoans: are reference genomes necessary to eliminate endosymbiont contamination and make robust phylogeographic inference? bioRxiv 2018; doi: https://doi.org/10.1101/440289

Tursch B, Tursch A. The soft coral community on a sheltered reef quadrat at Laing Island Papua New Guinea. Mar. Biol. 1982;68:321–332

Van Alstyne KL, Wylie CR, Paul VJ. Antipredator defenses in tropical Pacific soft corals Coelenterata: Alcyonacea II. The relative importance of chemical and structural defenses in three species of Sinularia. J. Mar. Biol. Ecol. 1994;178:17–34

van Oppen MV, Willis BL, Vugt HV, Miller DJ. Examination of species boundaries in the Acropora cervicornis group (Scleractinia, Cnidaria) using nuclear DNA sequence analyses. Mol Ecol. 2000;9:1363–1373

van Oppen MJ, McDonald BJ, Willis B, Miller DJ. The evolutionary history of the coral genus Acropora (Scleractinia, Cnidaria) based on a mitochondrial and a nuclear marker: reticulation incomplete lineage sorting or morphological convergence? Mol. Biol. Evol. 2001;18:1315–1329

van Oppen MJ, Willis BL, Van Rheede T, Miller DJ. Spawning times reproductive compatibilities and genetic structuring in the Acropora aspera group: evidence for natural hybridization and semi‐permeable species boundaries in corals. Mol. Ecol. 2002;11:1363–1376

Vargas S, Breedy O, Siles F, Guzman HM. How many kinds of sclerite? Towards a morphometric classification of gorgoniid microskeletal components. Micron. 2010;41:158–164

Verseveldt J. A revision of the genus Sinularia May Octocorallia Alcyonacea. Zoologische Verhandelingen. 1980;179:1–128

Vollmer SV, Palumbi SR. Hybridization and the evolution of reef coral diversity. Science. 2002;296:2023–2025

Willis BL, Babcock RC, Harrison PL, Wallace CC. Experimental hybridization and breeding incompatibilities within the mating systems of mass spawning reef corals Coral Reefs. 1997;16:S53–S65

Willis BL, van Oppen MJ, Miller DJ, Vollmer SV, Ayre DJ. The role of hybridization in the evolution of reef corals. Annu. Rev. Ecol. Evol. Syst. 2006;37:489–517

WoRMS Editorial Board. World Register of Marine Species. 2018 http://wwwmarinespeciesorg at VLIZ. Accessed 19 Dec 2018, doi:1014284/170

Wylie CR, Paul VJ. Chemical defenses in three species of Sinularia Coelenterata Alcyonacea: effects against generalist predators and the butterflyfish Chaetodon unimaculatus Bloch. J. Mar. Biol. Ecol. 1989;129: 141–160

Zwickl DJ. GARLI: genetic algorithm for rapid likelihood inference. 2006. http://www.bio.utexas.edu/faculty/antisense/garli/Garli.html, accessed 15 Jan 2019

