## Supplementary material for "A next generation approach to species delimitation reveals the role of hybridization in a cryptic species complex of corals": Quattrini_Supplfig1_Final.pdf

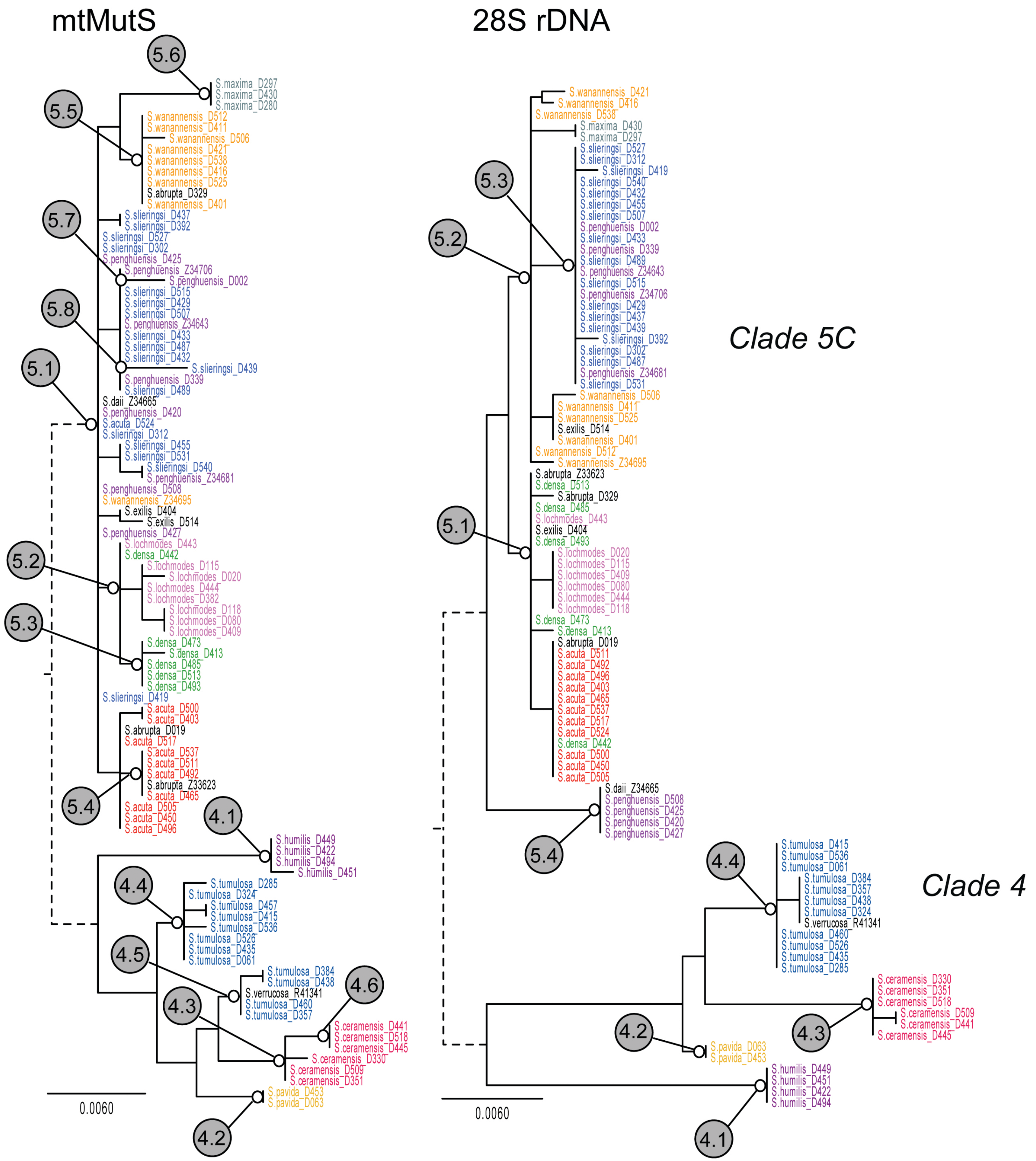


Maximum likelihood trees of barcode sequences for species in Sinularia clades 4 and 5C. (a) 735-bp mtMutS mitochondrial barcode. (b) 764-bp 28S rDNA nuclear barcode. Numbered circles indicate molecular operational taxonomic units (MOTUs) delimited using a 0.3% genetic distance threshold. Different morphospecies are color-coded. Trees are mid-point rooted with the branch connecting clades 4 and 5C pruned to facilitate readability.
