## Supplementary material for "A next generation approach to species delimitation reveals the role of hybridization in a cryptic species complex of corals": Quattrini_SupplFig2_BIC_Kclusters.pdf

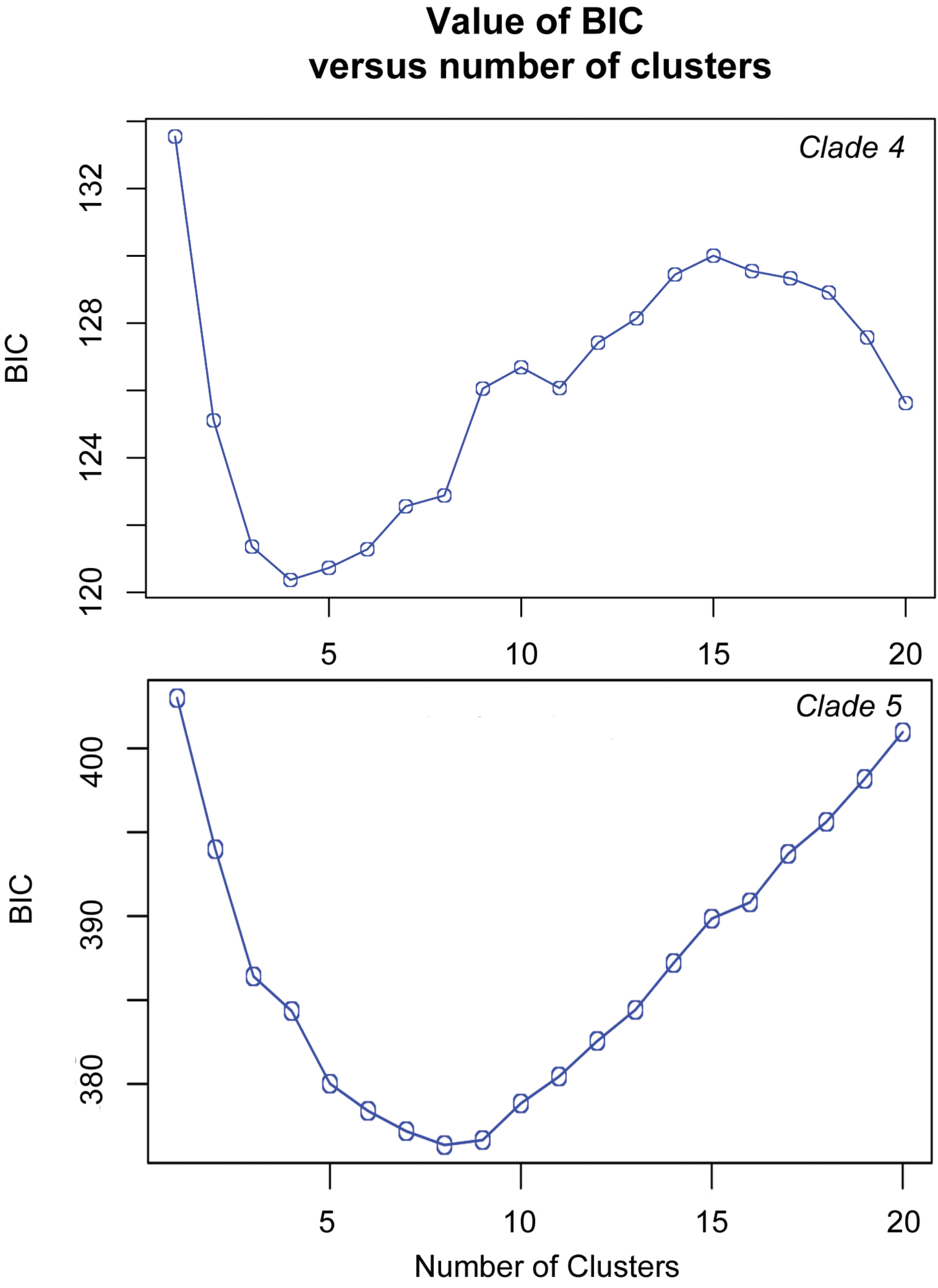


Bayesian Information Criterion (BIC) indicating the optimal number of k cluster. The lowest BIC often is determinant of the best number of k clusters.
