## Supplementary material for "A next generation approach to species delimitation reveals the role of hybridization in a cryptic species complex of corals": Quattrini_SupplFig3_clade4assignplot.pdf

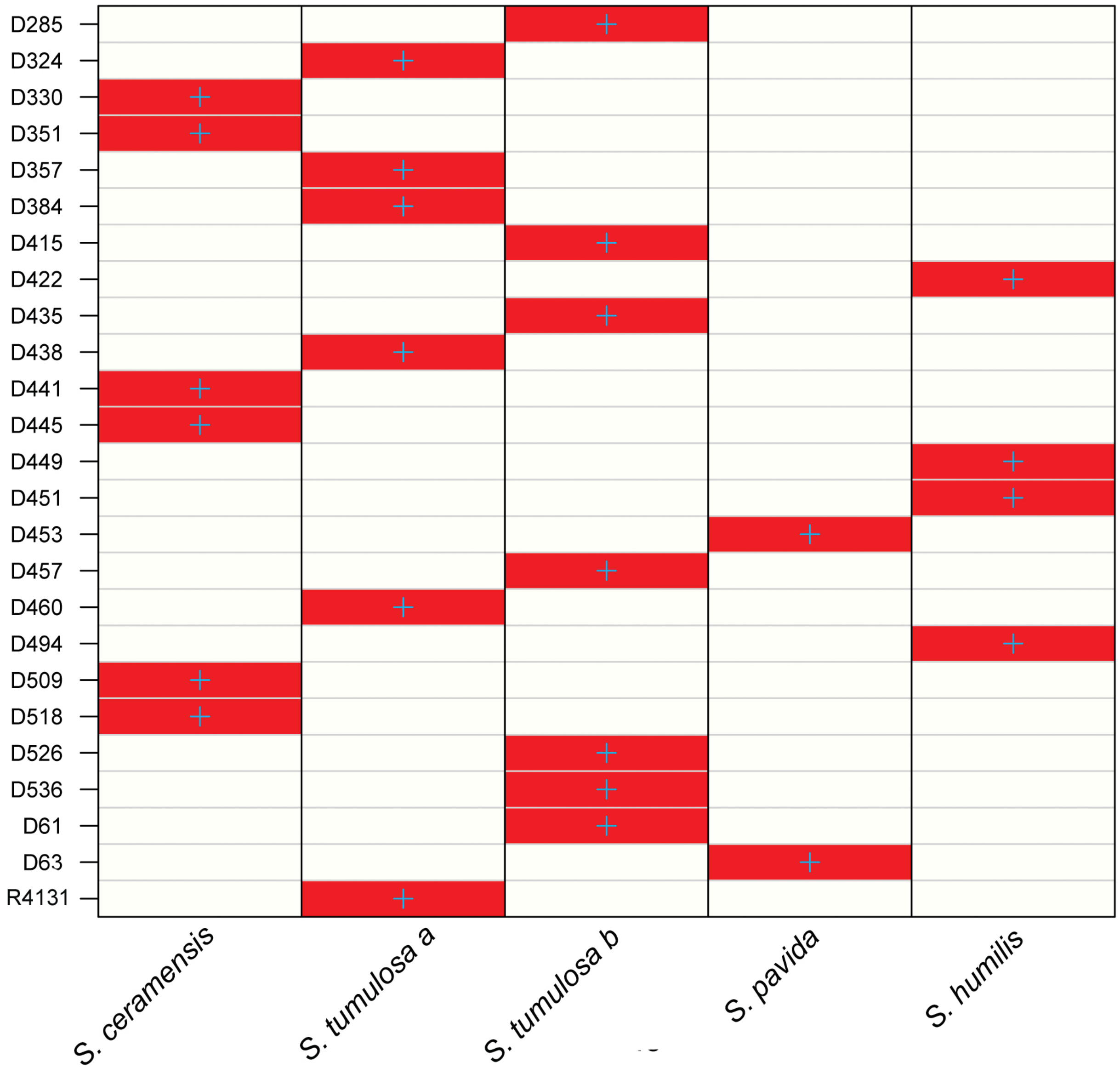


Probability of membership of each sample to a particular morphospecies assignment Successful reassignment is based on the discriminant functions of individuals to their original clusters.. Red=membership probability of 1.0. Blue crosses represent the prior cluster provided to DAPC
