## Supplementary material for "A next generation approach to species delimitation reveals the role of hybridization in a cryptic species complex of corals": Quattrini_SupplFig4_clade5assignplot.pdf

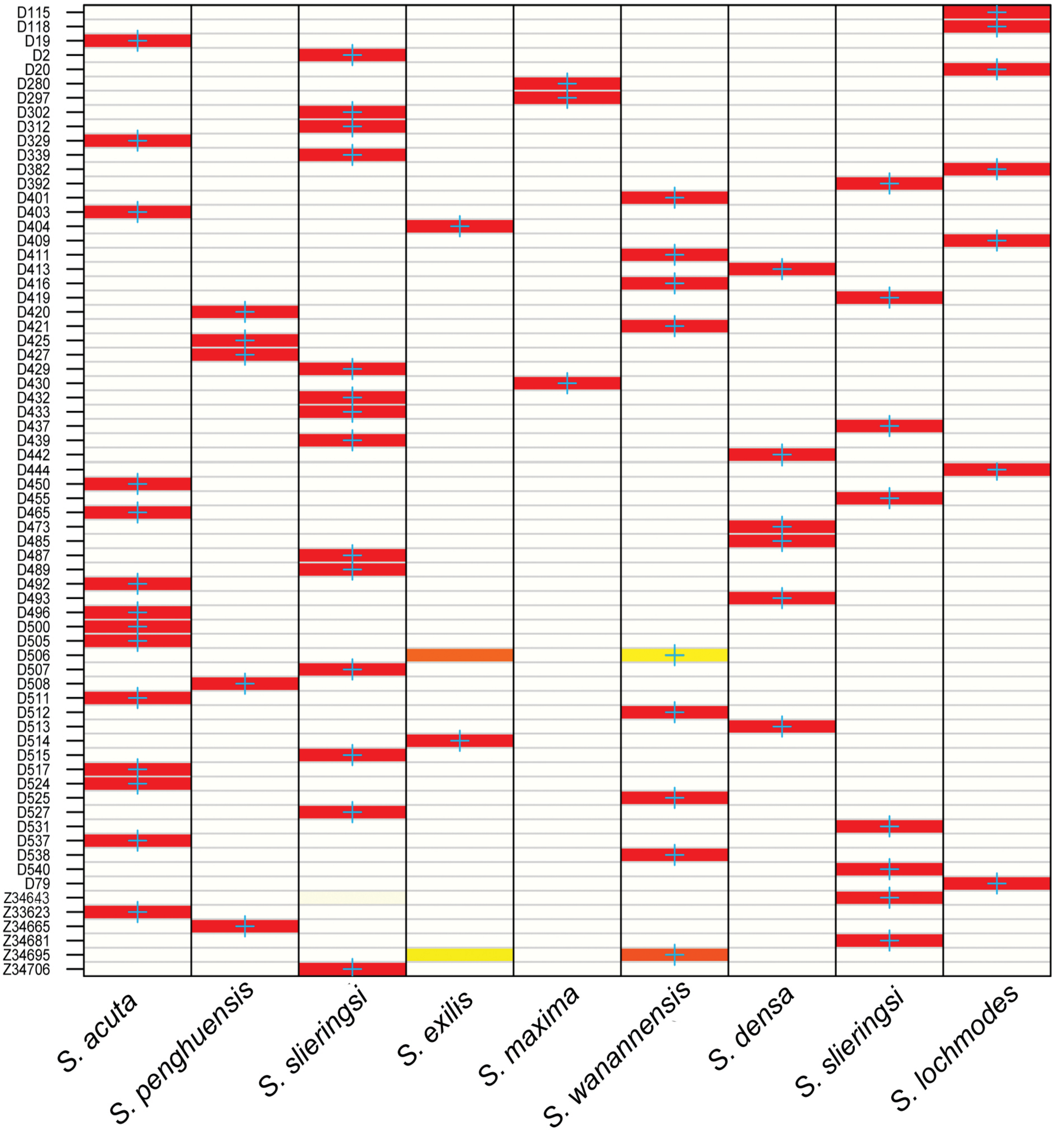


Probability of membership of each sample to a particular morphospecies assignment Successful reassignment is based on the discriminant functions of individuals to their original clusters. (Membership probability of 1=red,0= white. Blue crosses represent  the prior cluster provided to DAPC
