## Supplementary material for "A next generation approach to species delimitation reveals the role of hybridization in a cryptic species complex of corals": Quattrini_SupplFig5.pdf

**Clade 4** — mutS

| Clade 4 | M156 | M168 | M179 | M200 | M204 | M217 | M253 | M264 | M294 | M307 | M343 | M365 | M414 | M429 | M443 | M504 | M547 | M583 | M630 | M631 | M666 | M716 | M726 |
|---|---|---|---|---|---|---|---|---|---|---|---|---|---|---|---|---|---|---|---|---|---|---|---|
| 4.1 humilis | G | A | G | **C** | C | G | **A** | **C** | T | C | G | G | **C** | **C** | C | C | **A** | **T** | **T** | T | **A** | T | **T** |
| 4.2 pavida | G | G | G | T | C | G | G | T | T | C | G | **A** | T | T | C | **T** | G | C | C | **C** | G | T | C |
| 4.3 ceramensis | **A** | G | **R** | T | **T** | **A** | G | T | **C** | C | G | G | T | T | **M** | C | G | C | C | T | G | T | C |
| 4.4a tumulosa a | G | R | G | T | C | **A** | G | T | T | C | **A** | G | T | T | C | C | G | C | C | T | G | T | C |
| 4.4b tumulosa b | G | A | G | T | C | G | G | T | T | **Y** | G | G | T | T | C | C | G | C | **T** | T | G | **C** | C |

**Clade 4** — 28S

| Clade 4 | S073 | S076 | S106 | S109 | S122 | S126 | S171 | S185 | S206 | S225 | S232 | S289 | S315 | S362 | S363 | S365 | S366 | S447 | S454 | S515 | S516 | S530 | S531 | S532 | S533 | S534 | S535 | S536 | S537 | S538 | S539 | S540 | S541 | S542 | S543 | S544 | S545 | S551 |
|---|---|---|---|---|---|---|---|---|---|---|---|---|---|---|---|---|---|---|---|---|---|---|---|---|---|---|---|---|---|---|---|---|---|---|---|---|---|---|
| 4.1 humilis | **C** | **G** | A | **G** | G | **T** | **T** | G | C | **C** | **G** | **G** | C | C | A | G | **C** | **A** | **C** | A | **T** | **G** | **C** | **G** | **C** | **T** | **T** | **C** | **G** | **T** | **C** | **T** | **C** | **G** | **A** | C | G | C |
| 4.2 pavida | T | A | A | C | G | C | G | G | C | T | A | T | C | C | A | G | G | G | G | A | C | d | d | d | d | d | d | d | d | d | d | d | d | d | d | C | **A** | C |
| 4.3 ceramensis | T | A | A | C | **A** | C | G | **C** | C | T | A | T | **T** | C | **G** | **A** | **A** | G | G | **G** | C | d | d | d | d | d | d | d | d | d | d | d | d | d | d | C | G | T |
| 4.4a tumulosa a | T | A | **G** | C | G | C | G | G | **T** | T | A | T | C | **Y** | A | G | G | G | G | A | C | d | d | d | d | d | d | d | d | d | d | d | d | d | d | **T** | G | T |
| 4.4b tumulosa b | T | A | **G** | C | G | C | G | G | **T** | T | A | T | C | C | A | G | G | G | G | A | C | d | d | d | d | d | d | d | d | d | d | d | d | d | d | **T** | G | T |

**Clade 5C** — mutS

| Clade 5C | M144 | M151 | M224 | M245 | M248 | M249 | M285 | M547 | M597 | M618 | M621 | M634 | M690 |
|---|---|---|---|---|---|---|---|---|---|---|---|---|---|
| 5.1 daii/penghuensis | C | G | C | T | C | T | A | A | A | C | C | A | T |
| 5.2a slieringsi a | C | G | C | T | C | T | A | A | **M** | C | C | A | T |
| 5.2b slieringsi b | C | **R** | C | T | C | T | A | A | A | C | C | A | T |
| 5.3 acuta/abrupta | C | G | C | T | C | T | A | A | A | C | Y | A | T |
| 5.4 maxima | C | G | **T** | T | C | T | **C** | A | A | **T** | **T** | A | **C** |
| 5.5 densa | C | G | C | T | **A** | T | A | **T** | A | C | C | A | T |
| 5.6 wanannensis | C | G | C | T | C | **C** | A | A | A | C | **T** | A | T |
| 5.7 exilis | **T** | G | C | T | C | T | A | A | A | C | C | A | T |
| 5.8 lochmodes | C | G | C | **C** | C | T | A | **T** | A | C | C | **W** | T |

**Clade 5C** — 28S

| Clade 5C | S056 | S060 | S120 | S155 | S157 | S164 | S232 | S292 | S342 | S386 | S509 | S531 | S534 | S544 | S546 |
|---|---|---|---|---|---|---|---|---|---|---|---|---|---|---|---|
| 5.1 daii/penghuensis | G | G | **C** | **T** | **G** | **A** | **A** | **C** | G | G | C | **T** | C | C | **T** |
| 5.2a slieringsi a | G | **T** | T | C | C | d | G | T | **A** | G | C | C | C | C | C |
| 5.2b slieringsi b | G | **T** | T | C | C | d | G | T | **A** | G | C | C | C | C | C |
| 5.3 acuta/abrupta | G | G | T | C | **G** | d | G | T | G | G | **A** | C | **G** | C | C |
| 5.4 maxima | G | G | T | C | C | d | **A** | T | G | G | C | C | C | **G** | C |
| 5.5 densa | G | G | T | C | **G** | d | G | T | G | G | **A** | C | C | C | C |
| 5.6 wanannensis | K | G | T | C | C | d | R | T | G | **S** | C | C | C | C | C |
| 5.7 exilis | **T** | G | T | C | C | d | G | T | G | G | C | C | C | C | C |
| 5.8 lochmodes | G | G | T | C | **G** | d | G | T | G | G | **A** | **T** | C | C | C |

Suppl. Fig. X: Character-based barcodes for clade 4 and 5C *Sinularia* based on *mtMutS* and 28S rDNA sequences. Species are numbered to reflect the clade to which they were assigned in RADseq species delimitation analyses. Shaded, bolded nucleotides are species-diagnostic characters; bolded IUPAC ambiguity codes indicate polymorphic positions at which the alternative allele is private to a species while those shown in yellow are simple (non-diagnostic) polymorphisms. Each string of characters represents a "pure diagnostic combination" (deSalle et al. 2005) unique to a species. Nucleotide positions are numbered from the start codon of mtMutS and from the conserved motif AAACCGTTG in 28S (approx. 40 bp downstream of forward PCR primer 28S-Far).
