## Supplementary material for "A next generation approach to species delimitation reveals the role of hybridization in a cryptic species complex of corals": Quattrini_SupplFig6_optimAscore.pdf

### a-score optimisation - spline interpolation

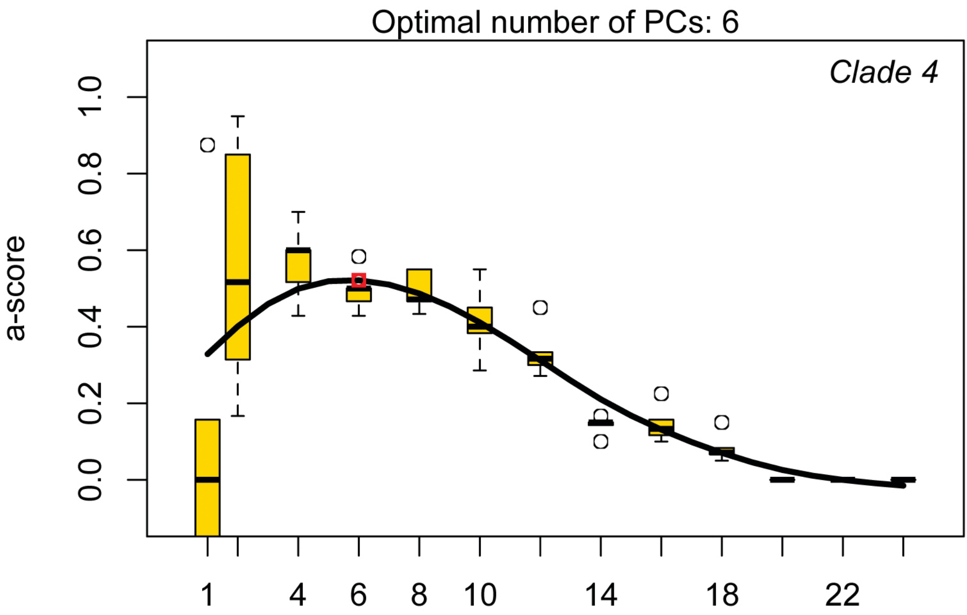


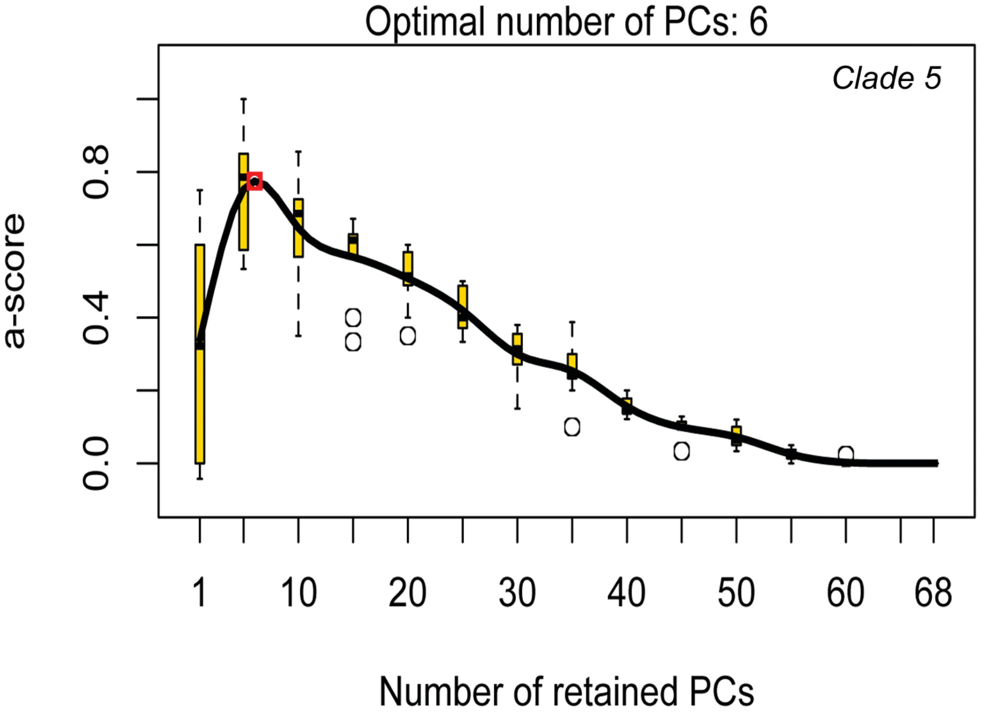


a-scores obtained from optim.a.score funciton in the R package adegenet, indicating (by red dot) the optimal number of principal components retained for both clades 4 and 5C.
