## Supplementary material for "A next generation approach to species delimitation reveals the role of hybridization in a cryptic species complex of corals": Quattrini_SupplFig9_clade5.spmodels.pdf

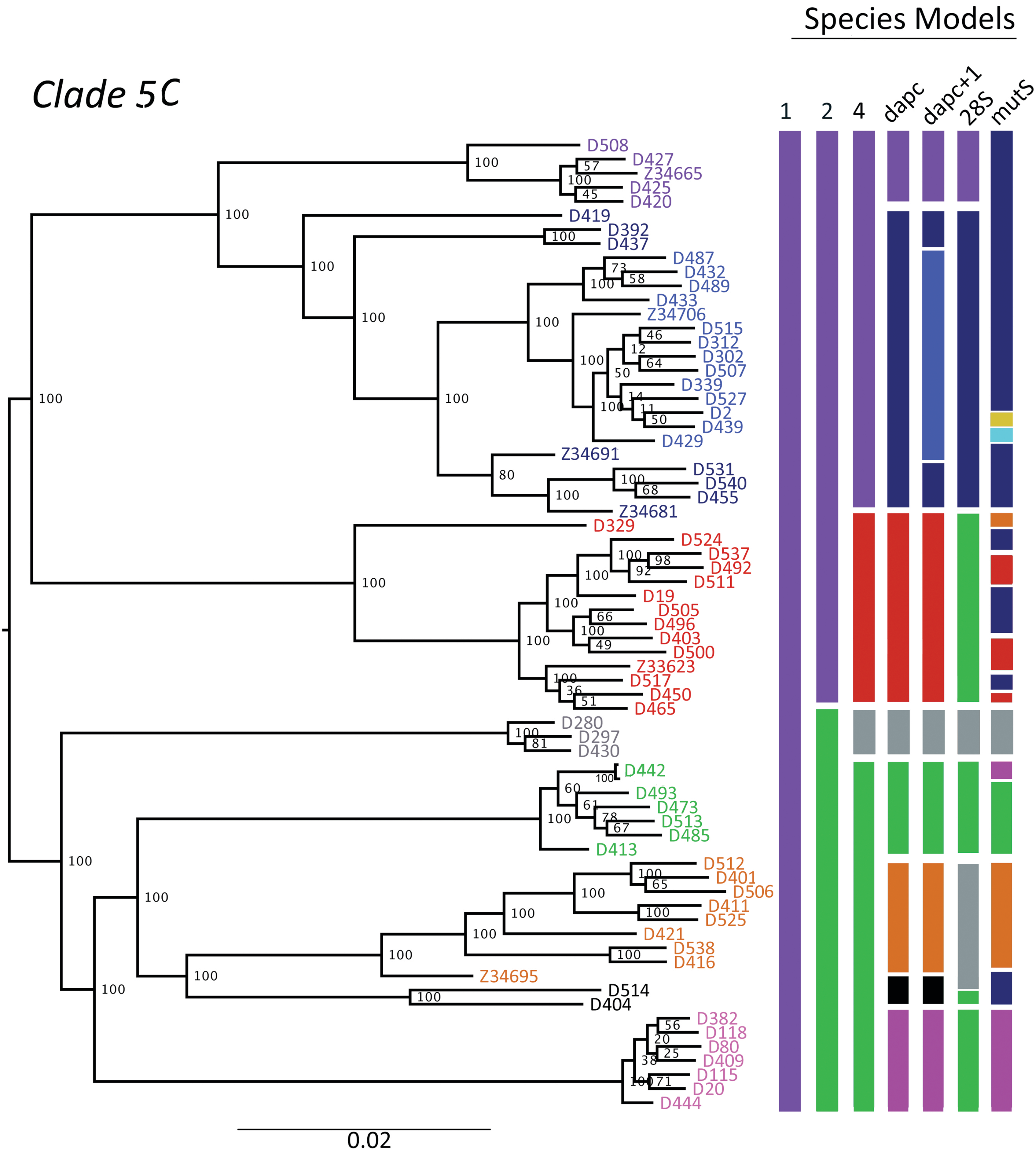


RAxML tree based on concatenated loci built with the m.75,c.85,p.25 dataset. Sample numbers are indicated. Species are color coded to match the species delimitations. Species models listed on the right were included in the BFD* analyses
