## Supplementary material for "A next generation approach to species delimitation reveals the role of hybridization in a cryptic species complex of corals": Quattrini_SupplFile5.docx

*RADSeq Analyses*

Sequence reads were de-multiplexed and trimmed for low quality using the *process_radtags* program in Stacks v 1.35 (Catchen et al., 2013). Illumina adapters and barcodes were excluded from reads, and lengths were truncated (-t) to 91bp. The rescue barcodes and RAD-tags option was enabled (-r) while reads with ambiguous bases (-c) and low quality scores (-q, <10) were discarded. The default sliding window (-w 0.15) was used with the default average score (-s 10) for filtering.

pyRAD v3.0 (Eaton, 2014) was then used for additional filtering and clustering of sequences to identify homologous loci (following Pante et al., 2015; Herrera & Shank 2016). We chose this program over other programs that also analyze RADSeq data because pyRAD enables the identification of homologous loci across highly divergent samples by including indel variation in the clustering process (Eaton, 2014). The maximum number of low quality (phred quality score <20) bases retained in each read was set at 5 and reads with more than 5 N’s and short reads (<50 bp) were discarded. Reads were then clustered at two different clustering thresholds (*c* 0.85, 0.90), which is the similarity threshold to cluster sequences within and across individuals. At a clustering threshold of *c* 0.85, we evaluated the minimum depth of coverage per cluster to make a base call (*d* 4, 6). Because few differences were found in the number of loci obtained between these two coverage depths, we restricted the depth of cluster to 6 for all pyRAD runs. The number of SNPs allowed in a final locus was set at 20. All pyRAD analyses were conducted on the entire *Sinularia* dataset (i.e., clades 4 and 5 together), but different combinations of datasets for clade 4 and clade 5C were output at the final step (step 7) in the pyRAD workflow. At this step, additional combinations of parameters were set at different clustering thresholds (*c* 0.85, 0.90): the minimum number of individuals per locus (*m* 0.50, 0.75, 1.0) and the maximum number of shared polymorphic sites (or shared heterozygous sites) at a locus (*p* 0.10, 0.25). Also at step 7, one individual (D446) was removed from the analysis because it was not in either clade 4 or 5.

The standalone version of DeconSeq v 0.4.3 (Schmieder & Edwards, 2011) was used to look for potential contamination by algal symbionts*,* Symbiodiniaceae. For each clade, loci were screened against the *Fugacium kawagutii* (formerly *Symbiodinium,* LaJeunesse et al., 2018) genome (Lin et al., 2015) (<http://web.malab.cn/symka_new/>). Less than 0.5% of the loci for each clade were found to be potential Symbiodiniaceae loci. Because so few Symbiodiniaceae loci were found and no differences were evident in the tree topologies (see methods below) with and without their inclusion, all potential Symbiodiniaceae loci were retained in the datasets for ease of bioinformatic analyses. Notably, a recent study demonstrated that clustering algorithms, such as those used by pyRAD, act as *de facto* filters for Symbiodiniaceae loci from datasets (Titus & Daly, 2018).

*Maximum Likelihood Phylogenies From Different RAD Datasets*

Topologies of the phylogenetic trees produced by different RADSeq locus datasets were similar for both clades 4 and 5C, particularly at the deeper nodes; however, some individuals moved around within their species-clade assignments. Four monophyletic clades matching morphospecies in clade 4 were well supported (100% b.s. support) across all ML phylogenies generated with different locus datasets (see Suppl. File 1 and Table 2). Several nodes, however, were poorly supported (<90% b.s. support) in the clade 4 phylogenies constructed with datasets that included no missing data (*m* 1.0, 73 to 185 loci per dataset, Table 2). Except for trees constructed from datasets with no missing data (*m* 1.0, 115-154 loci per dataset, Table 2), eight monophyletic clades corresponding to eight morphospecies were well supported (100% b.s. support) in all clade 5C ML phylogenies (see Suppl. File 1). Topologies were mostly congruent, except one individual, D002, moved from the *S. slieringsi* clade to the *S. lochmodes* clade when more missing data were included (*m* 0.50, 13,189 to 23,946 loci per dataset, Table 2). The clade 5C phylogenies generated with no missing data (*m* 1.0) were poorly supported (0-72% b.s support) and the eight monophyletic clades seen in the other phylogenies were largely absent. In addition, there were several unresolved polytomies in these phylogenies and many individuals of particular morphospecies shifted positions, grouping with other morphospecies in these datasets generated from few loci.

For each clade, the number of loci substantially decreased when allowing for 25% missing taxa per locus (>2,000 loci per individual) to no missing taxa per lous (<200 loci per individual). The substantial decrease in locus capture was likely due to the quality of DNA submitted for library preparation and sequencing. Most of the samples exhibited considerable DNA degradation, which can impact the number of loci recovered. Because some level of degradation is often unavoidable in field-collected material and in museum-based collections, perhaps future work should include more sequencing to increase coverage; the latter approach has been show to increase the phylogenetic utility of RADseq datasets (Eaton et al., 2017).
