## Supplementary figures and images for "A next generation approach to species delimitation reveals the role of hybridization in a cryptic species complex of corals"

### Quattrini_SupplFig7_DAE.pdf

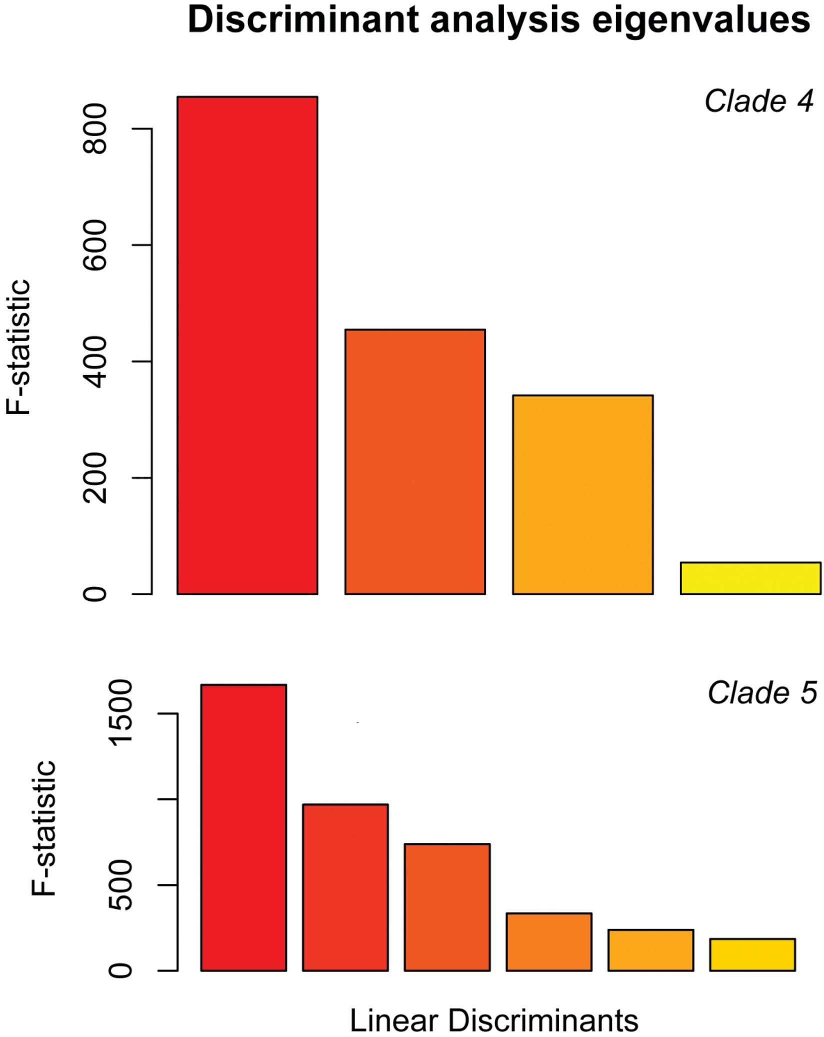


Total number of discriminant eigenvalues obtained in for DAPC analyses of clades 4 and 5C.
